# Integrated transcriptomic and proteomic analyses identify novel biomarkers of bladder outlet obstruction

**DOI:** 10.64898/2026.04.29.721732

**Authors:** Alexander A. Bigger-Allen, Barnali Das, Yang Tang, Kyle Costa, Gabriel Luis Ocampo, Ali Hashemi Gheinani, Shannon DiMartino, Jane Kaull, John W. Froehlich, Richard S. Lee, Rosalyn M. Adam

**Affiliations:** Urological Diseases Research Center, Boston Children’s Hospital, Boston, MA, USA; Department of Surgery, Harvard Medical School, Boston, MA, USA; Functional Urology Research Group, Department for BioMedical Research DBMR, University of Bern, Switzerland; Department of Urology, Inselspital University Hospital, 3010 Bern, Switzerland

**Keywords:** Spinal cord injury, neurogenic bladder, urine biomarkers, inosine, multi-omics

## Abstract

Bladder outlet obstruction leads to pathological remodeling and emergence of lower urinary tract symptoms. Although relief of obstruction is associated with symptomatic improvement, it is not universally successful, reflecting persistent alterations in the bladder. Reliable surrogate biomarkers of obstruction are lacking, particularly early in the disease course before irreversible damage to the bladder may have occurred. In this study, re-analysis of publicly available transcriptomic datasets from diverse rodent models of obstruction identified tissue transcripts including *Cthrc1*, *Grem1*, *Ltbp2, Acta1* and *Msn* that were induced in response to injury. Candidate markers were validated experimentally in an independent model of neurogenic obstruction demonstrating time-dependent changes, and a subset (*GREM1*, *LTBP2*, *ACTA1*, *MSN*) was increased in bladder tissue from pediatric patients with neurogenic bladder. Several candidates including CTHRC1, ACTA1 and MSN were also differentially abundant in independent proteomic datasets from a rat model of partial bladder outlet obstruction. Candidate markers were also attenuated in datasets from rats with either surgical removal of obstruction or treatment with anticholinergic medication or inosine. Integrated analysis of tissue transcriptomics data and tissue and urine proteomics data from a model of neurogenic obstruction revealed modest but statistically significant concordance between markers observed in tissue and urine. These findings identify tissue and urine biomarkers of both non-neurogenic and neurogenic obstruction that may reflect early changes in obstructive uropathy that could be monitored in a non-invasive manner.

## Introduction

Bladder outlet obstruction arises from both anatomical causes such as prostatic enlargement and pelvic organ prolapse, as well as neurogenic causes, including spinal cord injury. Irrespective of the etiology, both anatomic and functional obstruction share common pathways of pathological bladder wall remodeling and the emergence of lower urinary tract symptoms (LUTS)(1). Although the obstruction can be removed in some patients, LUTS are not always relieved completely, which may be related to persistent bladder alterations (reviewed in (2)). Monitoring the longitudinal evolution of clinically meaningful obstruction, i.e. that which evokes structural and functional changes within the bladder, is challenging since existing methods such as urodynamics are invasive and impractical in asymptomatic patients. However, once symptoms occur, irreversible damage to the bladder may have occurred. Reliable surrogate biomarkers of bladder deterioration would enable longitudinal monitoring of bladder health and, in combination with symptom improvement, would also have the potential to measure response to intervention. A particular unmet need is for non-invasive biomarkers that detect early, biologically meaningful bladder remodeling before irreversible dysfunction occurs.

Bladder wall remodeling in response to obstruction of both neurogenic and non-neurogenic origin reflects cellular changes that arise to compensate for increased outlet resistance and intravesical pressure, and include altered metabolism, perturbed blood flow, proliferation and extracellular matrix turnover. Sustained pressure increases, however, drive inflammation, fibrosis, bladder deterioration and functional decline (3, 4). Different analytes in urine have been explored as potential biomarkers in the context of outlet obstruction. These include growth factors, enzymes, metabolites and microRNAs that are associated with the fibroproliferative remodeling that results from obstruction, whether neurogenic or non-neurogenic. Among these, urinary levels of the neurotrophins NGF and BDNF, ATP, MMP-1 and-2, TGFβ1, prostaglandin E2, markers of oxidative stress and discrete miRNAs have been linked to bladder dysfunction in obstructed patients (5–13). In addition, several markers including NGF and BDNF have also demonstrated sensitivity to treatment (14, 15), although some studies have raised doubts over NGF as a reliable marker for neurogenic dysfunction (16). Despite these associations however, NGF, BDNF and markers of oxidative stress are also elevated in cystitis and bladder pain syndromes (17–24), and therefore lack specificity for outlet obstruction.

An ongoing question related to outlet obstruction is defining the earliest point at which functionally significant pathological alterations in the bladder are evident and how these evolve over time (reviewed in (4)). While evaluation of urine is non-invasive, the extent to which changes in urinary analytes reflect local bladder remodeling rather than systemic or renal contributions remains incompletely defined, as are the similarities and differences between markers of non-neurogenic and neurogenic obstruction. Given these challenges, the potential utility of biomarkers to monitor bladder health in the setting of obstruction has not yet translated reliably into clinical practice (25).

Based on shared pathophysiology characterized by fibroproliferative remodeling, we hypothesized that conserved molecular responses across diverse models of obstruction would identify robust biomarkers that are detectable in urine and reflective of underlying alterations in bladder tissue. In this study, we have leveraged publicly available gene expression data from rodent models of bladder outlet obstruction to assess (i) acute and chronic transcriptional signatures evoked by obstruction; (ii) the extent to which markers are shared in bladder outlet obstruction of diverse etiologies; (iii) how marker expression changes over time after injury; (iv) how markers change in response to surgical or pharmacological intervention; and (v) the concordance between tissue transcriptome, proteome and urinary proteome to gauge the extent to which markers present in urine reflect alterations in bladder tissue. We have validated a subset of these findings experimentally in rat tissue and urine in the setting of neurogenic obstruction.

## Results

### Early gene signatures of bladder outlet obstruction

To identify gene expression patterns that reflect the response of the bladder to outlet obstruction, we re-analyzed 4 publicly available datasets from different injury models in the rat, including: (i) partial urethral ligation (26); (ii) pelvic ganglion denervation (27); (iii) obstruction secondary to suprasacral spinal cord transection (SCT) (28); and (iv) our own rat SCT dataset from a recent study (29). To capture the acute response, we restricted the analysis to the earliest time period common to all models, using expression profiles obtained at 10-14 d following injury (**Table 1, Suppl. Fig. 1A-D**). All datasets comprised transcriptomic data from full-thickness rat bladder tissue. Interestingly, despite these datasets sharing the same species, tissue of interest and comparable time period, the differences are extensive and highlighted in **Suppl. Fig. 2A-C**. As a result, we hypothesized that shared changes would be biologically meaningful and suggest common indicators or mechanisms of early stages of obstruction regardless of etiology. Accordingly, we prioritized genes consistently altered across models as candidate markers of core, early responses of the bladder to obstruction, as opposed to model-specific changes.

**Table 1.**
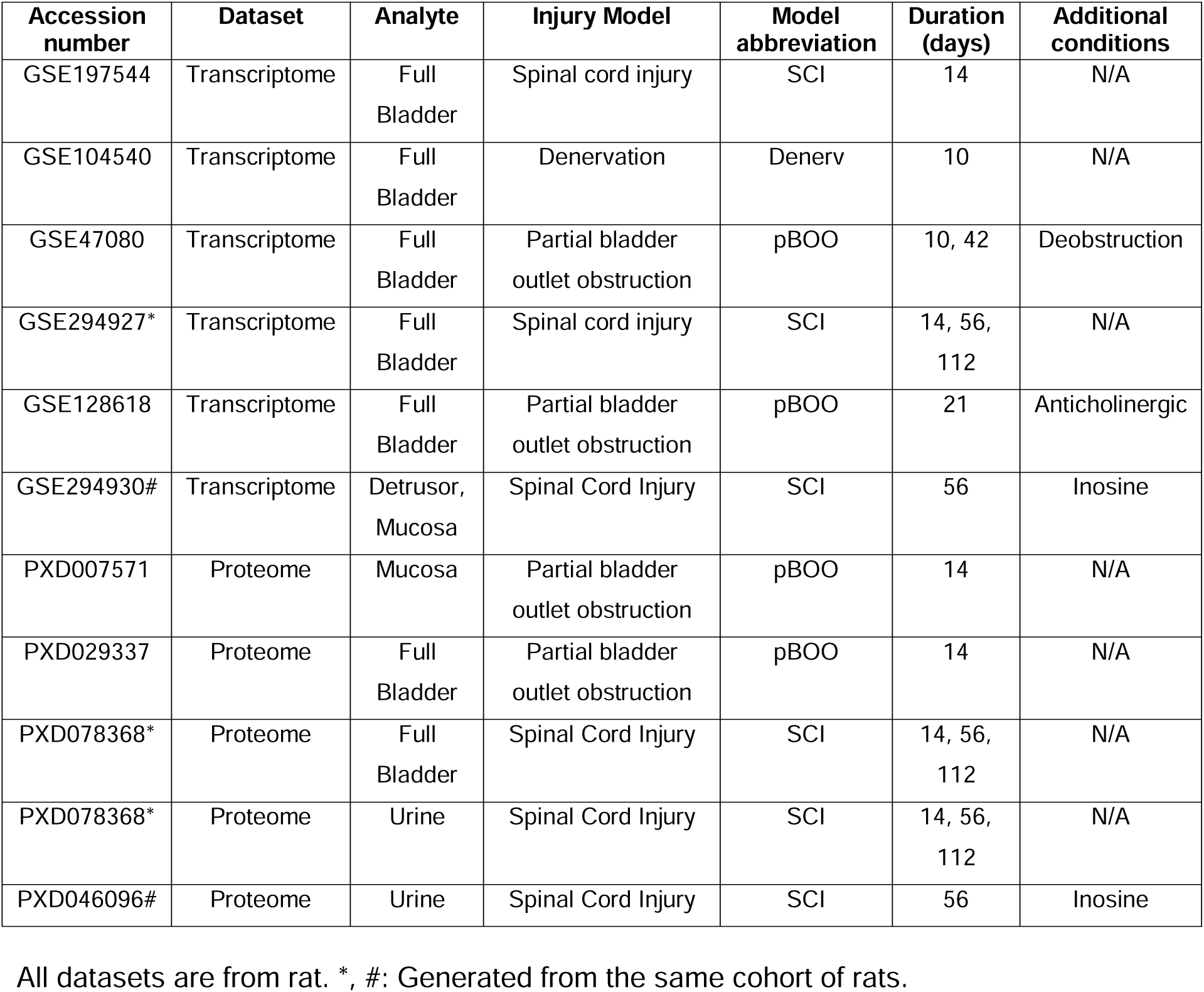
Public datasets from NCBI GEO and ProteomeXchange.

Comparative analysis identified 44 differentially expressed genes (DEGs) across all 4 datasets that were altered in common by injury in comparison to the respective non-injured controls. The shared DEGs in all 4 datasets included 11 that were upregulated (**Fig. 1A**) and 33 that were downregulated (**Fig. 1B**). To identify which pathways were in common and unique across different models of obstruction, we performed enrichment analysis based on GO terms and KEGG pathways. Among the most enriched pathways, those associated with DNA damage, actin cytoskeleton regulation, cardiomyopathy, and inflammation were shared across injury models (**Fig. 1C**), consistent with our previous findings, which identified pathological changes related to DNA damage, contractility, cytoskeletal components and ECM-receptor interactions in the bladder following spinal cord injury (29). Notably, the most commonly occurring DEGs associated with these pathways included *Itgb1*, *Acta1*, *Col4a1*, *Col4a2*, *Pak1*, *Mylk*, *Tmod1* and *Tnnt2*, emphasizing the role of cytoskeletal and contractile changes as central processes dysregulated by obstruction. Together, these observations reflect the hallmarks of obstruction-induced remodeling, namely oxidative stress and DNA damage, integrin-mediated mechanotransduction, smooth muscle hypertrophy, extracellular matrix deposition and altered contractility.

**Figure 1:**
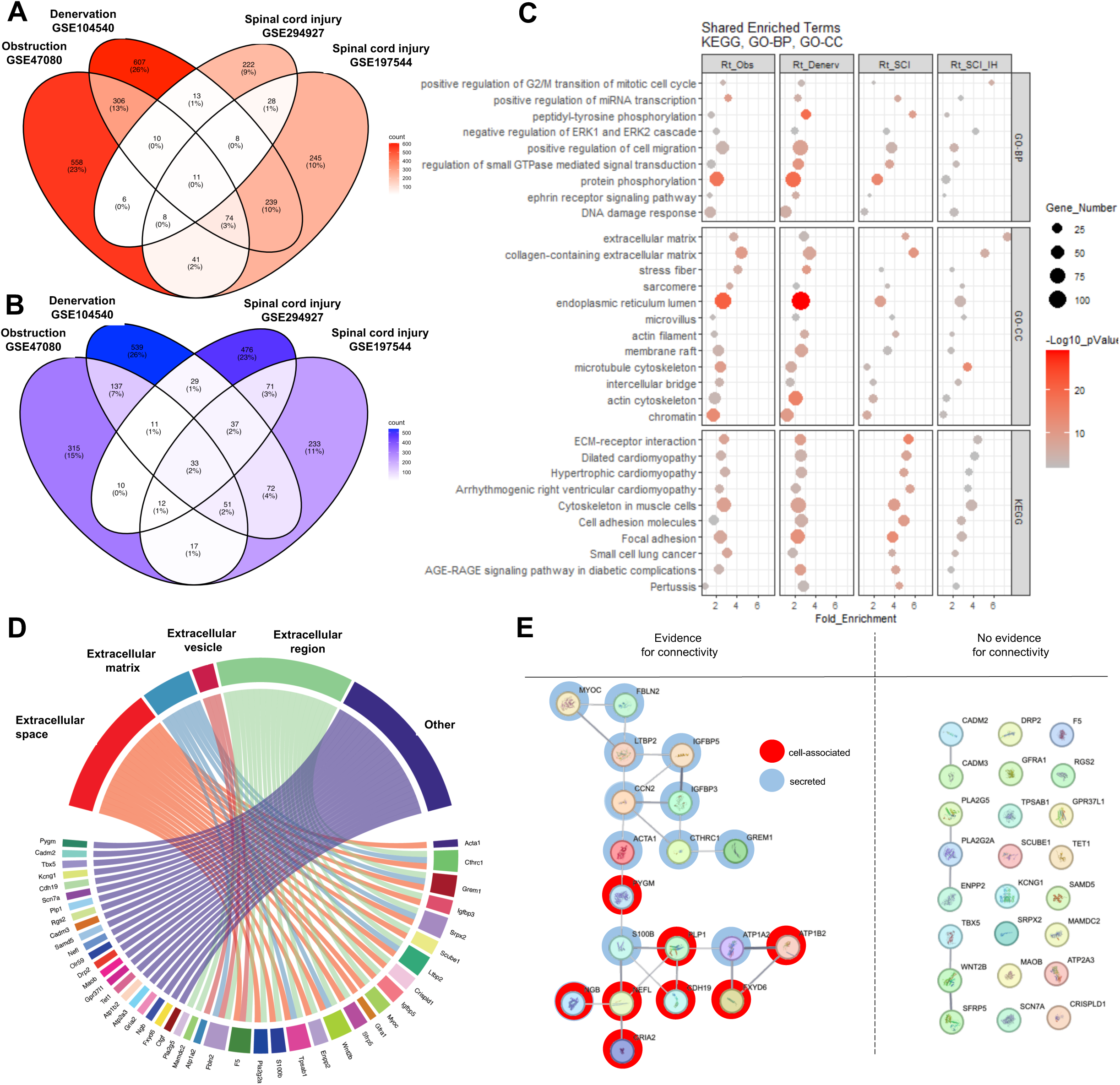
Early gene signatures of rodent models of bladder outlet obstruction. Differentially expressed genes (DEGs) from all four rat models of bladder injury were compared using Venn diagrams. Up-regulated **(A)** and down regulated (**B)** DEGs are colored in increasing intensities of red and blue, respectively, proportional to the number of genes in each comparison. **(C)** Dot plot of clustered enrichment analysis of shared KEGG pathways and GO-terms related to Biological Process and Cellular Compartment using all DEGs for each dataset. The extent of enrichment is on the x-axis and the enriched pathways are on the left y-axis. Pathways enriched in each dataset were subjected to clustering analysis and are organized into clusters on the right y-axis. The size of the dots indicates the number of genes associated with a given pathway from the list of DEGs of each dataset. The color intensity of the dots indicates the statistical significance of the enriched pathway. **(D)** Chord diagram of the shared DEGs from all 4 datasets are connected to GO terms related to cellular compartments associated with the extracellular space. Color coding is arbitrary and enhances contrast between the features. **(E)** STRING network analysis identified 19 potential biomarkers that have evidence for contributing to a shared function or pathway. These biomarkers were highlighted with colored circles based on up-regulation (red) or down-regulation (blue) in all datasets. Biomarkers with no evidence for connectivity, having an edge confidence value of less than 0.4 were not included in the network assembly.

Next, to identify markers which may be secreted and detectable in body fluids, thereby enriching for candidates with potential translational utility as non-invasive urinary biomarkers, we filtered the 44 shared DEGs responsive to injury using gene ontology (GO) terms related to cellular compartment such as extracellular space, extracellular matrix, exosome and extracellular vesicle, or none of these (**Fig. 1D**). Of these, 22 were defined as extracellular and 22 were associated with another cellular compartment. For those DEGs defined as extracellular we determined potential interactions among them or contributions to the same biological process using STRING network analysis. As shown in **Fig. 1E**, STRING analysis identified 19 targets with evidence of contributing to a shared process or pathway and of these, 11 (*Acta1, Cthrc1, Grem1, Igfbp3, Ltbp2, Igfbp5, Myoc, Fbln2, Atp1a2, S100b, Ccn2*) were predicted to be extracellular based on GO terms.

### Time-dependent changes in early gene signatures of bladder outlet obstruction

To understand temporal changes in early markers of obstruction, we explored expression of a subset of the shared DEGs over time after injury in a single model. We chose our model of functional obstruction secondary to spinal cord injury since it spans acute (2 weeks), established (8 weeks) and chronic (16 weeks) time points (29). In this analysis, transcriptomic data were generated from full-thickness bladder tissue. Using these data, we identified DEGs showing statistically significant differences in expression at any of the three time points following SCI compared to uninjured controls **(Fig. 2A)**. Visualization of the counts for early, secreted (**Fig. 2B)** and cell-associated (**Fig. 2C)** biomarkers predicted from GO-term association and STRING network analysis from **Fig. 1D & E** revealed distinct patterns of expression over time. Notably *Acta1*, *Fbln2*, *Cthrc1*, and *Ltbp2* emerged as transcripts showing expression patterns in tissue from injured versus uninjured animals that were distinct between time points. This contrasts with the cell-associated biomarkers which show uniform up-or down-regulation across all time points. These data suggest that the expression of ‘secreted/extracellular’ biomarkers can discriminate uniquely between the time points induced by injury compared to controls, whereas cell-associated biomarkers may be limited to identifying changes induced by injury without temporal resolution. To determine the extent to which DEGs associated with obstruction could discriminate SCI from controls, we performed principal components analysis using all features (**Fig. 2D**), the 44 DEGs shared across different injury models (**Fig. 2E**), the 11 secreted (**Fig. 2F**), and the 8 cell-associated biomarkers (**Fig. 2G**). Based on this analysis, the 11 secreted biomarkers enabled discrimination of early and established/chronic SCI from all control time points. These findings suggest that extracellular biomarkers may provide temporal resolution of disease progression, in contrast to cell-associated markers which primarily reflect the presence of injury.

**Figure 2.**
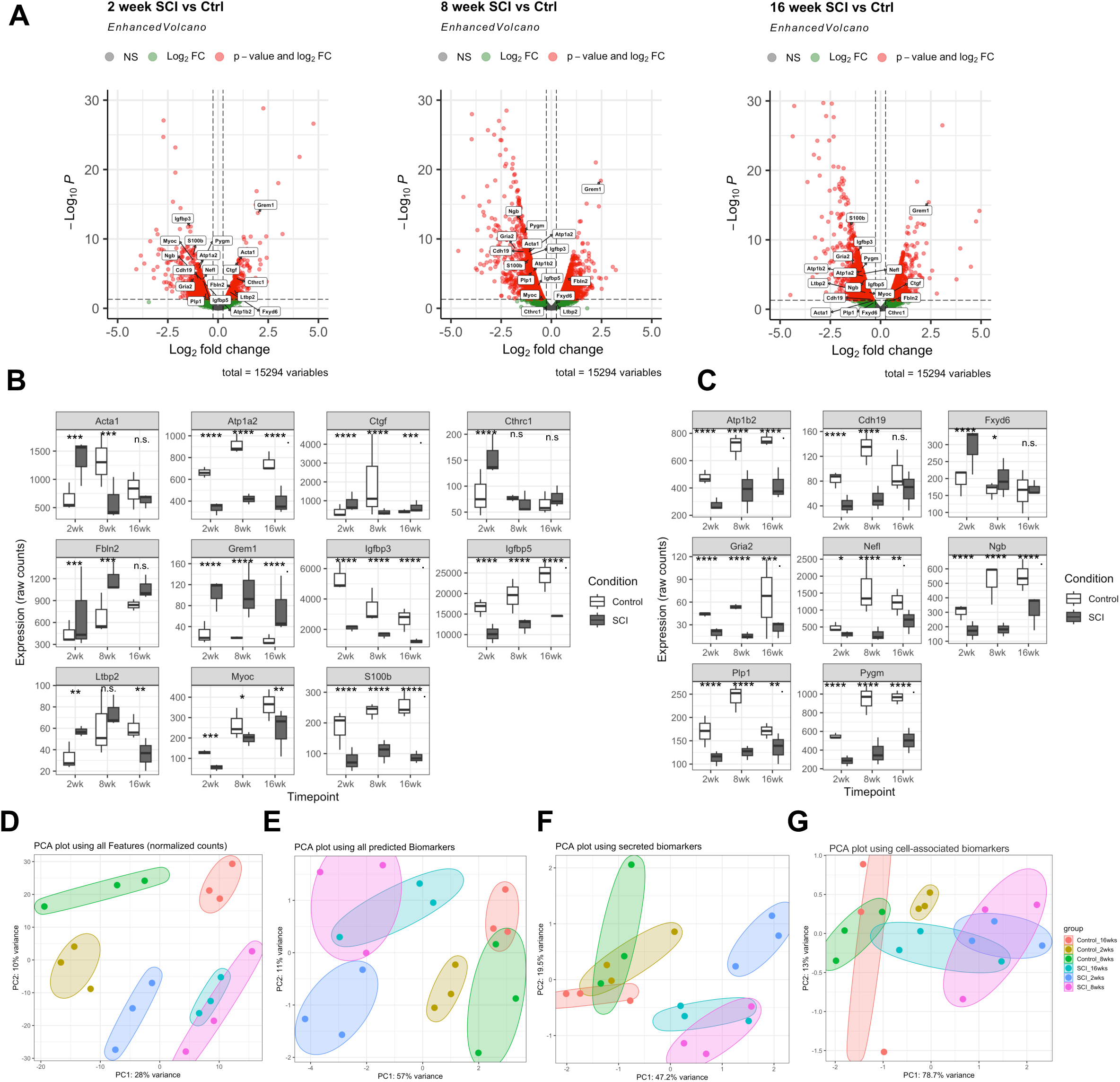
Predicted soluble biomarkers persist in chronic phase of injury and distinguish SCI injured bladders from controls. **(A)** Volcano plots labeled with the 11 shared, up-regulated DEGs predicted to be biomarkers at each time point of spinal cord injury relative to uninjured controls. **(B)** Boxplots of raw counts for the 11 shared, extracellular biomarkers and **(C)** 8 cell associated biomarkers (n = 3 replicates). PCA plot of all replicates for each condition using the entire count matrix of features **(D)**, all 19 predicted biomarkers **(E),** the 11 extracellular candidate biomarkers **(F)** or the 8 cell-associated biomarkers **(G)**.

### Predicted biomarkers are sensitive to treatment of obstruction

To further filter the list of predicted biomarkers, we assessed their sensitivity to treatment of obstruction. Two public datasets were leveraged to answer this question. The first public dataset, GSE47080, comprised transcriptomic data from full-thickness bladder tissue from rats subjected to partial bladder outlet obstruction for i) 10 days, ii) 6 weeks, or iii) 6 weeks followed by 10 days of de-obstruction and was re-analyzed for patterns in which genes were differentially expressed at 6 weeks after pBOO and attenuated with surgical relief of obstruction (**Fig.3A).**

**Figure 3.**
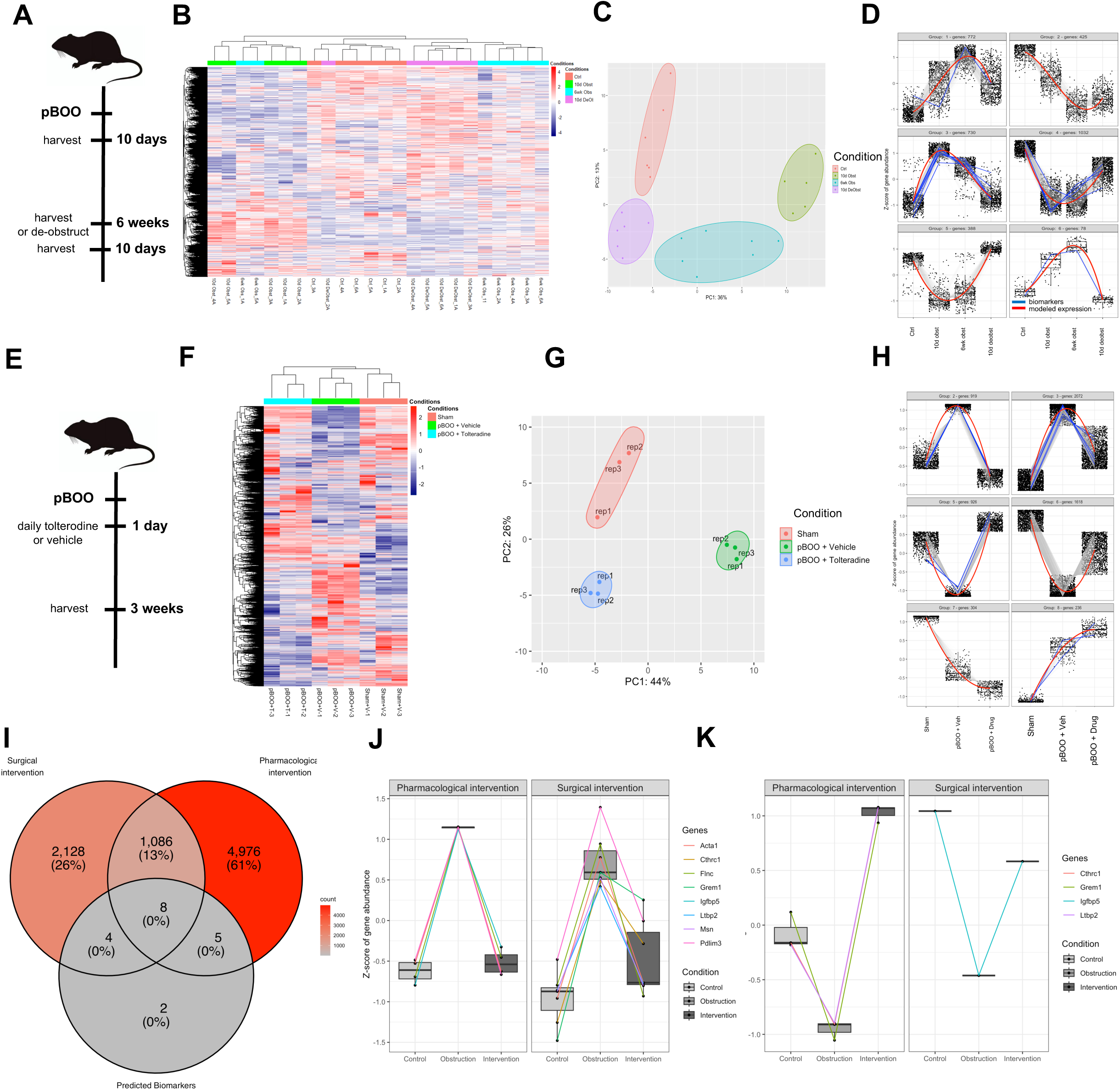
Biomarkers of bladder obstruction are sensitive to intervention. (**A)** Schematic of pBOO time course and conditions used in GSE47080. **(B)** Heatmap of clustering analysis of log10-transformed gene expression of the most variable genes; color coding is scaled by row. **(C)** Principal component analysis of all 4 groups in the dataset. (**D)** Pattern analysis of ∼3500 differentially expressed genes in bladders of rats subjected to 10d obstruction (10d Obst), 6wk obstruction(6wk Obst), 6wk obstruction followed by 10d of de-obstruction (10d DeObst) versus controls (Ctrl) highlighting patterns of genes sensitive to injury and surgical intervention. Red lines highlight the modeled gene expression pattern across all genes for a given group. Blue lines highlight the expression of predicted biomarkers. **(E)** Schematic of pBOO time course and conditions used in GSE128618. **(F)** Heatmap of clustering analysis of log10 transformed gene expression of the most variable genes; color coding is scaled by row. **(G)** Principal component analysis of all 3 groups in the dataset. **(H)** Pattern analysis of ∼6000 differentially expressed genes in bladders of rats subjected to 3wks obstruction (pBOO), obstruction + tolterodine (pBOO + tolterodine) versus controls (Sham) highlighting patterns of genes sensitive to pBOO in the absence and presence of anticholinergic treatment. Red lines highlight the modeled gene expression pattern across all genes for a given group. Blue lines highlight the expression of predicted biomarkers. **(I)** Venn diagram showing intersection of 19 predicted biomarkers and differentially expressed genes sensitive to obstruction and intervention in both datasets. **(J)** Box plots of z scores of biomarker gene abundance increased by injury and attenuated by pharmacological or surgical intervention. **(K)** Box plots of z scores of biomarker gene abundance decreased by injury and attenuated by pharmacological and surgical intervention.

Clustering analysis of the most variably expressed genes showed that samples from de-obstructed rats clustered between control samples and those following 6 weeks of obstruction **(Fig. 3B).** A similar pattern was observed with principal component analysis, with de-obstructed samples clustering more closely with control samples than with obstructed samples, reflecting resolution, at least in part, of obstruction-induced gene expression changes with surgical intervention **(Fig.3C).** Next, we performed pattern analysis on ∼3500 genes that were differentially expressed across all groups (10 d Obst, 6 wk Obst, 6 wk Obst/10 d De-Obst) relative to controls (Ctrl) to identify genes that were sensitive to both obstruction and de-obstruction. Among these, genes that were upregulated with obstruction (Group 3) and genes that were downregulated with obstruction (Group 4), which together comprised nearly half of the DEGs, were returned to near baseline following removal of obstruction **(Fig. 3D)**, suggesting these genes are dynamically regulated in response to obstruction and its relief.

A second public dataset, GSE128618, was analyzed to determine biomarkers sensitive to pharmacological intervention (30). This dataset comprises transcriptomic data from full-thickness bladder tissue from rats subjected to pBOO for 3 weeks, in which the animals were treated daily with vehicle or the anticholinergic agent tolterodine **(Fig. 3E)**. Clustering analysis of the most variably expressed genes clustered the replicates of each group together **(Fig. 3F)**.

Principal component analysis showed a loss in variation between the control and tolterodine treated pBOO bladders in the first principal component **(Fig. 3G),** suggesting that gene expression changes that contributed to differences between pBOO and control were reduced and attributable to the effect of tolterodine. To identify tolterodine-sensitive DEGs, we performed pattern analysis and identified 4 gene expression patterns (**Fig. 3H)**: genes up-or down-regulated with obstruction that were partially or completely reversed with tolterodine treatment. Between the de-obstruction and tolterodine datasets, nearly 1100 DEGs were shared and among them were 8 of the 19 predicted biomarkers **(Fig. 3I)** described earlier (Fig.1E and Fig.2D). These genes sensitive to both pharmacological and surgical intervention included *Acta1*, *Cthrc1*, *Flnc*, *Grem1*, *Igfbp5*, *Ltbp2*, *Msn*, and *Pdlim3* **(Fig. 3J, K)**. Among these intervention-sensitive DEGs only Msn was associated with any of the top 20 pathways dysregulated by pBOO in both datasets **(Suppl. Fig. 3A-B)**.

### Multi-omics comparison of time-dependent changes in bladder tissue and urine following SCI

Given the limited concordance between the transcriptome and proteome reported in the literature (31, 32) we next assessed the extent of correlation between mRNA and protein levels in the setting of obstruction by performing a comparative analysis of the bladder tissue transcriptome, bladder tissue proteome, and the urine proteome from our rat SCI time course. The time points included 2, 8 and 16 weeks after SCI which represent early, established, and chronic phases of experimental neurogenic bladder, respectively. The subsequent analyses were performed using either i) the 161 targets differentially expressed in all three datasets at all three time points **(Fig. 4A)** or ii) using >6,800 differentially expressed targets identified in any of the three analytes at any time point **(Suppl. Fig. 4A)**. In the focused analysis of the 161 shared DEGs/differentially expressed proteins (DEPs), bladder transcriptome data at all time points cluster together with the 2-week urine proteome. In the comprehensive comparison, transcriptomic data at all time points cluster with the 2-and 8-week urine proteome. However, no time point of the bladder transcriptome clustered with the bladder tissue proteome. This was the case in both the restricted **(Fig. 4A)** and the comprehensive **(Suppl. Fig. 4A)** analyses.

**Figure 4.**
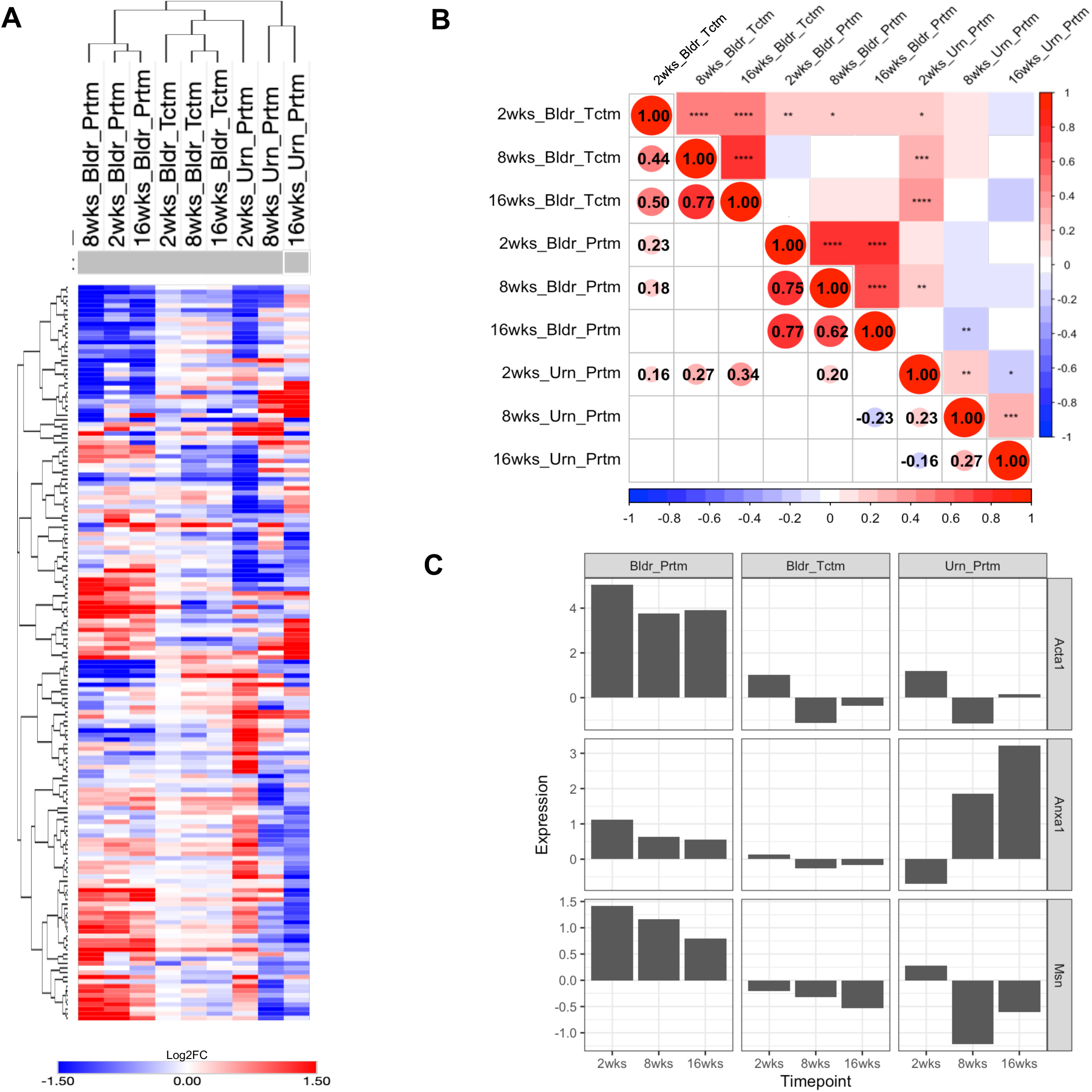
Multi-omic comparison of bladder tissue and urine over time following SCI in the rat model. **A)** Heatmap of the log2FC of all 161 targets shared between all datasets at all time points. Hierarchal clustering of rows and columns using 1-pearson correlation was employed to quantify the similarities between each dataset. **B)** Correlation plots quantify the extent to which each time point of each dataset correlates. Top half of correlation plot indicates the extent of statistical significance in the correlation of each time point and dataset. The bottom half explicitly shows the R^2^ value between the dataset that have statistically significant correlation. **C)** Bar chat showing average fold change expression of shared targets that are differentially expressed in multiple time points in all datasets.

Next, we performed pairwise correlation analysis to quantify (using Pearson correlation coefficient) correlation using only the shared targets between any two analytes and time points. The 2-week bladder transcriptome showed positive correlation with the 2-and 8-week bladder proteome (r = 0.23, pval < 0.01 and r = 0.18, pval < 0.05 respectively) **(Fig. 4B)**. This correlation was stronger in the comprehensive analysis (r = 0.49, pval < 0.01 and r = 0.62, pval < 0.01 respectively) **(Suppl. Fig. 4B)**. The DEGs and DEPs in bladder tissue at 16 weeks after SCI showed a positive correlation (r = 0.25, pval < 0.05) only in the comprehensive analysis **(Suppl. Fig. 4B)**.

We next assessed whether the limited correlation between the bladder transcriptome and the bladder proteome changed when separating bladder tissue into detrusor and mucosa prior to assessing the transcriptome. Using separate cohorts of SCI or control rats at 8 weeks after injury, we compared the DEGs from detrusor and mucosa (GSE294930) with DEGs from full thickness (FT) bladder (GSE294927) and DEPs from FT bladder (PXD078368)**(Suppl. Fig 5A**). We found the number of DEGs in FT tissue (n=3665) was comparable to the sum of DEGs in detrusor (n=2465) plus mucosa (n=919) minus the 241 targets in common, i.e. n=3143 **(Suppl. Fig. 5B)**. Thus, in terms of absolute number of DEGs identified, use of FT tissue did not diminish the yield compared to separated tissues. However, ∼75% of the DEGs identified in FT tissue were not shared with either the detrusor or the mucosa, indicating that tissue separation substantially influences which genes are called differentially expressed **(Suppl. Fig. 5D-F)**.

Further, correlation analysis of only the shared differentially expressed targets in any two datasets showed a greater Pearson correlation coefficient between the detrusor DEGs and the FT DEPs (r = 0.40, p < 0.001) compared to the FT DEGs and DEPs (r = 0.29, p < 0.001). The mucosa DEGs showed lower correlation with the FT bladder proteome compared to the FT tissue DEGs (r = 0.27, p < 0.01) **(Suppl. Fig. 5C)**. These data suggest that changes induced by SCI in the bladder proteome, are in part, better reflected by changes in the expression of the genes in the detrusor. This is consistent with the detrusor constituting the bulk of the bladder wall, such that the FT bladder proteome is dominated by detrusor-derived proteins.

Comparison of the enriched KEGG pathways across the four sets of differentially expressed targets from tissue **(Suppl. Fig. 6A, 6C)** revealed 233 (FT DEGs), 216 (detrusor DEGs), 117 (mucosa DEGs) and 55 (FT DEPs) pathways, of which 16 were shared across all datasets **(Suppl. Fig. 6B).** Nearly 90% of the enriched pathways from the FT DEPs were among the 233 pathways from the FT DEGs, with 83% of pathways from FT DEPs enriched when compared to pathways of detrusor DEGs, and ∼50% of mucosa KEGG pathways present in pathways from FT DEPs. Correlation between enriched KEGG pathways from the FT DEGs and FT DEPs showed statistically significant, positive correlation using the fold enrichment score (r = 0.31, p<0.01). However, the pair-wise comparison between pathways enriched in detrusor or mucosa with pathways of FT DEPs showed greater correlation when using fold enrichment score (r = 0.42, p< 0.0001 and r = 0.54, p< 0.01, respectively). These data support that the limited extent of overlap between DEGs and DEPs regardless of tissue separation likely reflects changes in different components of the same perturbed pathways.

Next, we compared DEGs and DEPs from tissues with the urinary DEPs following SCI at multiple time points. Interestingly, the DEPs in urine at 2 weeks after SCI showed increasing positive correlation and statistical significance in a time-dependent manner with the DEGs of the FT bladder following SCI (2 wks: r = 0.16, pval < 0.05, 8 wks: r = 0.27, pval < 0.001, and 16 wks: r = 0.34, pval < 0.0001) **(Fig. 4B)**. A similar trend was observed in the comprehensive analysis reaching statistical significance between 8 and 16 weeks following injury (r = 0.25, pval < 0.001 and r = 0.41, pval < 0.0001 respectively) **(Suppl. Fig. 4C)**. These findings suggest that changes in the urine proteome are concordant with, and may reflect alterations in the bladder tissue transcriptome induced by SCI. In addition to comparing the datasets and time points at the level of DEGs and DEPs, we performed enrichment analysis using KEGG pathways. Nine pathways were enriched across the nine combinations of time points and analytes, the top three of which were ‘Cytoskeleton in Muscle Cells’, ‘Motor Proteins’ and ‘Focal Adhesions’ **(Suppl. Fig. 4C)**. Extending the comparison between urine DEPs and DEGs from detrusor and mucosa, demonstrated the highest correlation with the Detrusor DEGs (r = 0.35, p < 0.001) **(Suppl. Fig. 5C)**. Calculating the Pearson correlation coefficient using relative gene regulation for each KEGG pathway revealed the strongest correlation between pathways enriched in urine and mucosa at 8 wks after SCI (r = 0.45, p < 0.01) **(Suppl. Fig. 6C)**. Notably, beyond the pair-wise correlation, 12 targets were associated with the 9 shared pathways identified across all datasets in at least 1 time point **(Suppl. Fig. 4D)**. Of these targets, only *Acta1, Anxa1* and *Msn* were identified as shared across all time points and datasets **(Fig. 4C)**.

To validate our findings in an independent model of bladder injury, we reanalyzed two separate publicly available proteomics datasets comprising mucosa (PXD007571)(33) and FT bladder tissue (PXD029337)(34) from rats subjected to pBOO for 2 weeks. Of the roughly 700 proteins shared across >= 50% of the mucosa samples, nearly 200 DEPs were identified of which a subset included 3 biomarkers identified in our existing analysis --ACTA1, MYL6, and GSMT1 **(Suppl. Fig. 7A-E).** Similarly, in the FT tissue proteome, roughly 4000 proteins were identified in >= 50% of the samples and of those, 5 biomarkers identified in our existing analysis were among the list of ∼600 DEPs with a |log2FC| > 0.5 **(Suppl. Fig. 8)**. These included MSN, MYL6, FLNA, FBLN2, and CTHRC1. These data further support the identification of shared targets at the mRNA and protein level in both neurogenic and non-neurogenic bladder obstruction that may serve as biomarkers. However, given the small number of biological replicates and the incomplete sample annotation in these publicly available datasets, these findings should be regarded as supportive rather than confirmatory.

### Validation of time-dependent, predicted biomarkers in rat and human neurogenic bladders

To determine the potential translational value of the markers identified above, we assessed their expression first in bladder tissue from rats with SCI-induced neurogenic bladder and then in detrusor and mucosa from pediatric patients with neurogenic bladder and controls as a proof-of-concept for translation to human subjects. Among the markers predicted across the different models of injury, *Cthrc1, Grem1*, *Ltbp2, Acta1* and *Msn* mRNA were differentially expressed in bladder tissue from rats with neurogenic bladder in 1 or more time points compared to uninjured controls **(Fig. 5A)**. In detrusor and mucosa tissue from pediatric patients with neurogenic bladder *GREM1, LTBP2, MSN and ACTA1* were significantly increased compared to controls in at least 1 sample as measured by qPCR **(Suppl. Fig. 9)**. Analysis of urine specimens from the rat SCI time course revealed increased levels of MSN in urine early after injury in the rat, that declined at 8 and 16 weeks, whereas GREM1 levels showed a biphasic response following SCI, peaking at 8 weeks and declining thereafter **(Fig. 5B, 5C)** in agreement with tissue levels. CTHRC1 levels did not differ significantly between samples from injured versus control animals and LTBP2 was undetectable in rat urine.

**Figure 5.**
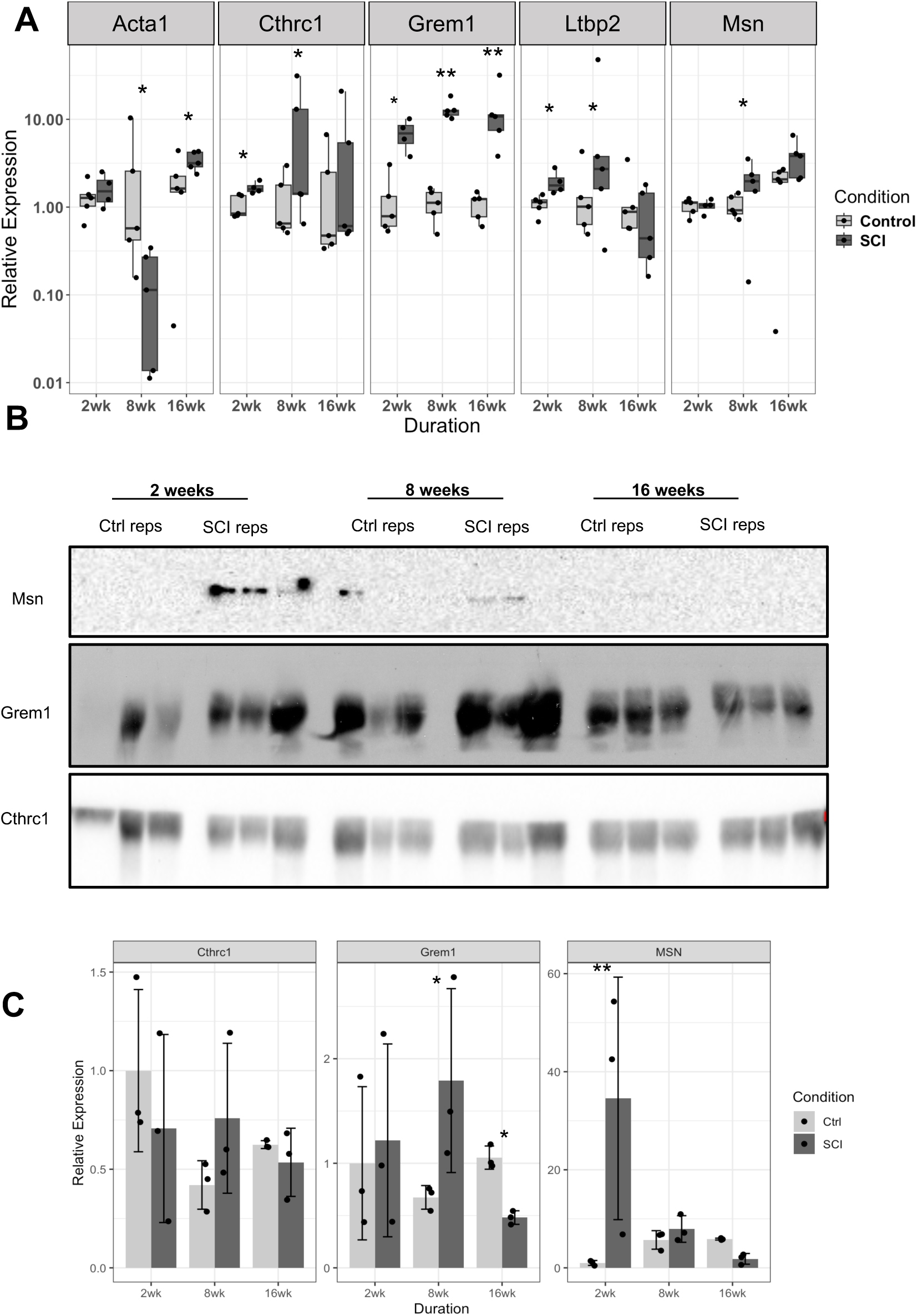
Validation of SCI induced gene and protein expression of predicted biomarkers. Validation of predicted biomarkers sensitive to intervention via **A)** qPCR of rat bladder tissue following 2, 8 and 16 weeks of spinal cord injury. n= 3 per condition and time point. **B)** Immunoblot of rat urine following 2, 8 and 16 weeks of SCI and controls. **C)** Quantification of bands in (B). Data were assessed for normality using Kruskal-Wallis and post-hoc analysis was a Dunn test to calculate p-values. * p-value < 0.05, ** p-value < 0.01, *** p-value < 0.001.

### Biomarkers in urine induced by SCI are attenuated by inosine

After identifying a subset of concordant, predicted biomarkers in the bladder transcriptome and urine proteome following SCI and pBOO in rodents, we sought to determine the extent to which these biomarkers were sensitive to pharmacological intervention with inosine. In previous studies we showed that inosine administration reduced detrusor overactivity in vivo in SCI rats (35), attenuated spontaneous activity in bladder tissue from SCI rats in vitro (36) and prevented pathological changes in the bladder induced by SCI, including reactive oxygen species, DNA damage, and pathways associated with aberrant contraction (29), suggesting that markers modulated by inosine are reflective of neurogenic bladder pathophysiology. In our SCI model in which rats received inosine daily for 8 weeks (29), many of the same pathways associated with SCI were enriched in the urinary DEPs compared to controls **(Suppl. Fig. 10A, B)**. Interestingly, 4 candidate biomarkers were among the urinary DEPs up-regulated in vehicle-treated SCI rats (SCI-Veh) compared to uninjured controls and were down-regulated in urine from inosine-treated SCI rats (SCI-Ino) compared to SCI-Veh **(Fig 6B, C)**. These included MSN, EZR, IGFBP3, and IGFBP5. Additionally, inosine not only downregulated these biomarkers in the context of SCI generally, but was able to reduce the abundance of MSN, EZR and IGFBP3 to 39%, 30% and 0.3% of control expression levels, respectively **(Fig. 6D)**. This suggests that these urinary proteins are associated with injury and sensitive to inosine treatment. In an unbiased assessment of all ∼1400 DEPs from across the three comparisons, we identified 1037 urinary DEPs that were dysregulated in SCI (SCI-Veh) and attenuated by inosine (SCI-Ino)**(Fig. 6E)**. Enrichment analysis using KEGG pathways based on the DEPs for i) SCI-Veh compared to control and ii) SCI-Inosine compared to SCI-Veh, showed several pathways identified in previous datasets of obstruction were sensitive to inosine **(Fig. 6F)** including ‘Focal adhesion’, ‘Regulation of actin cytoskeleton’, ‘TGF-beta signaling pathway’ and ‘Dilated cardiomyopathy’. Importantly, Msn and Ezr were among 16 inosine-sensitive urinary DEPs that were most highly associated with the top pathways dysregulated by SCI **(Fig. 6G, Suppl. Fig. 10A-B)**. A summary of the evidence supporting each candidate biomarker across all analyses is provided in Table 2.

**Figure 6.**
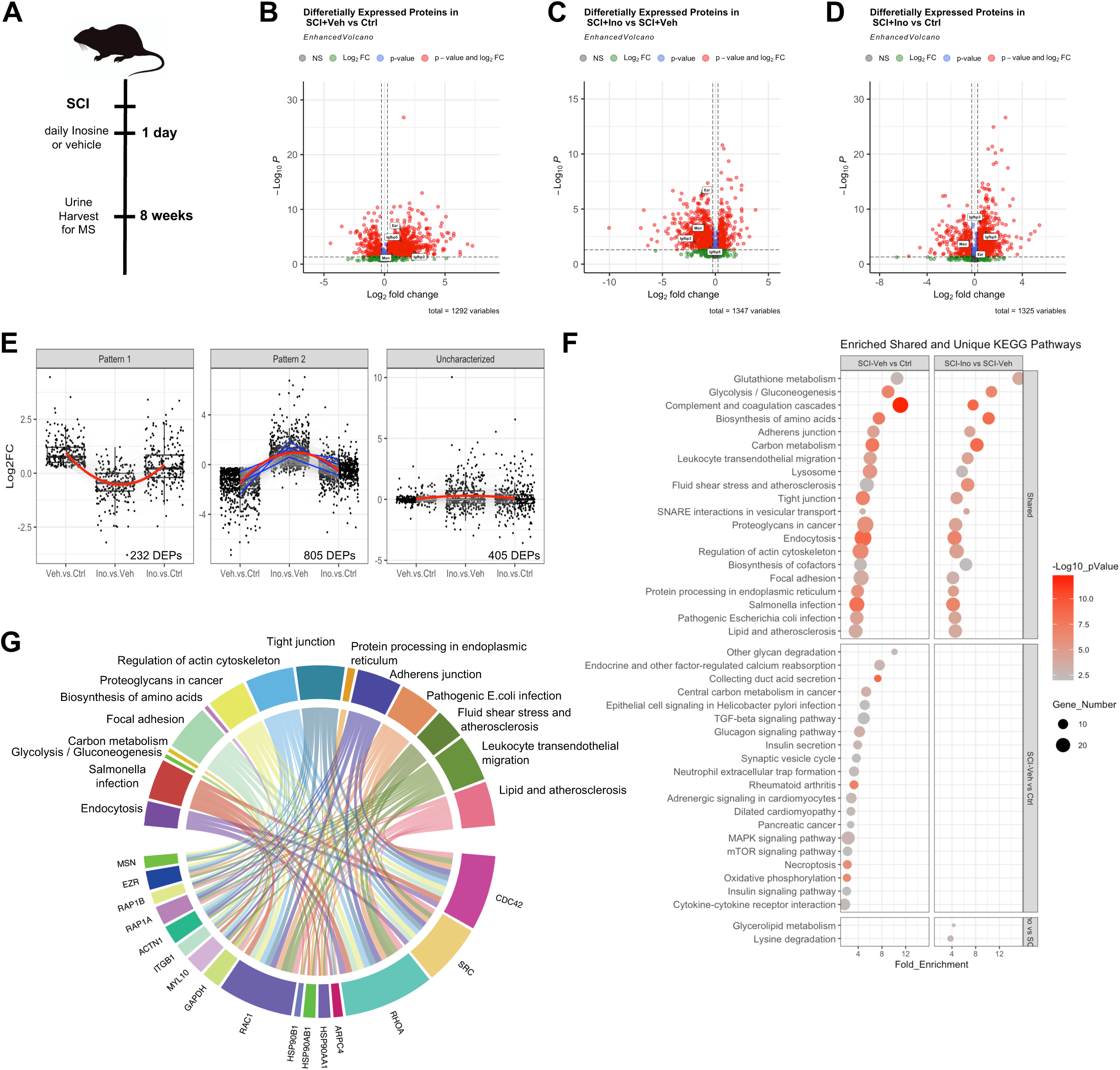
Inosine attenuates a subset of biomarkers in urine protein induced by SCI. **A)** Schematic of the duration and inosine intervention used in the spinal cord injury (SCI) rat model prior to urine collection and mass spectrometry analysis of urinary proteome. **B)** Volcano plot of the differentially expressed proteins (DEPs) in urine induced by SCI + vehicle vs. control (SCI-Veh vs Ctrl). **C)** Volcano plot of the differentially expressed proteins (DEPs) in urine induced by SCI + inosine vs. SCI-Veh (SCI-Ino vs SCI-Veh). **D)** Volcano plot of the differentially expressed proteins (DEPs) in urine induced by SCI + Inosine vs. control (SCI-Ino vs Ctrl). Red dots indicated DEPs that passed adjusted p-value < 0.05 and |log2FC| > 0.25. **E)** Boxplots depicting the patterns of inosine sensitive urinary DEPs based on log2FC in all 3 comparisons. **F)** Dotplot of enriched KEGG pathways using the statistically significant DEPs in pattern 2 of panel E) from the SCI-Veh vs. Ctrl and SCI-Ino vs. SCI-Veh comparisons. Dot size reflects the number of genes in each pathway, Dot color reflects the-log10 transformed p-value, the right y-axis indicates whether the pathway is shared or unique to across the two comparisons. All terms were enriched with a p-value <= 0.05. **D)** A chord diagram connecting the urinary DEPs to their associated pathways. Only DEPs associated with 4 or more pathways are shown.

**Table 2.** Summary of evidence for shared candidate biomarkers of bladder outlet obstruction.

| Candidate | Time-dependent (SCI) | Sensitive to intervention | Protein (pBOO) | DE in tissue and urine | Rat tissue qPCR | Rat urine immunoblot | Human tissue qPCR |
|---|---|---|---|---|---|---|---|
| **Acta1** | ✓ | ✓ | ✓ mucosa | ✓ | ✓ | n.t. | ✓ |
| **Cthrc1** | ✓ | ✓ | ✓ FT | – | ✓ | n.s. | n.s. |
| **Grem1** | ✓ | ✓ | – | – | ✓ | ✓ | ✓ |
| **Ltbp2** | ✓ | ✓ | – | – | ✓ | n.d. | ✓ |
| **Msn** | ✓ | ✓ | ✓ FT | ✓ | ✓ | ✓ (2 wk) | ✓ |

✓, supported; –, not identified; n.s., not significant; n.d., not detectable; n.t., not tested; FT, full thickness;

## Discussion

In this study we describe the identification and validation of candidate biomarkers of non-neurogenic and neurogenic bladder outlet obstruction in experimental models. Key aspects of the analysis include (i) evaluation of multiple independent datasets reflecting bladder tissue gene expression early after obstruction; (ii) investigation of candidate markers longitudinally up to 16 weeks after obstruction; (iii) integrated analysis of tissue transcriptome and proteome with urine proteome; (iv) assessment of candidate biomarker response to intervention; and (v) confirmation of a subset of candidates in bladder tissue from pediatric patients with neurogenic bladder. From this comprehensive analysis, Grem1, Cthrc1, Ltbp2, Acta1 and Msn emerged as biomarkers of bladder outlet obstruction that show persistent differential expression after obstruction, are sensitive to relief or treatment of obstruction, and for a subset (MSN, GREM1, ACTA1), are altered in urine in an experimental model of obstruction. Taken together, these findings suggest that urinary levels of MSN, GREM1 and ACTA1, together with tissue levels of CTHRC1 and LTBP2 may reflect biologically meaningful changes in the bladder associated with obstruction.

Strengths of our study include the incorporation of multiple independent datasets for candidate biomarker discovery and in different analytes, i.e., tissue mRNA, tissue protein and urinary protein. Consistent with prior reports, we observed modest concordance between the bladder tissue transcriptome and proteome following SCI with Pearson correlation coefficients of 0.2-0.6, depending on time point and feature set (32, 37, 38). In addition, we observed time-dependent, statistically significant concordance between the tissue transcriptome and the urine proteome suggesting that urine analysis may enable predictions about the underlying state of the tissue. Additional strengths include the validation of candidate markers at multiple time-points after injury and investigation of their response to intervention. While several urinary markers have been described previously in the setting of obstruction, these have typically been evaluated at a single timepoint during the disease course. Our study focused not only on the acute phase following obstruction, but also established and chronic phases thereby providing insights into the evolution of changes evoked in both tissue and urine in response to obstruction. Furthermore, interrogation of datasets in which obstruction was treated either surgically or pharmacologically identified genes that were responsive to relief of obstruction and those that remained differentially expressed despite treatment. The former group, i.e. those responsive to treatment in the setting of obstruction are informative with respect to disease burden, whereas those that did not change are candidates for driving persistent dysfunction in subjects with obstruction. Collectively, our study has not only identified candidate markers of disease burden and evolution, but also those sensitive to intervention.

Our analyses of separated detrusor and mucosa provide insight into the tissue sources of the changes captured by bulk bladder and urine analyses. Although the total number of DEGs identified in FT tissue was comparable to that obtained from the separated compartments, the majority of DEGs differed between FT and separated tissue, whereas concordance was considerably greater at the pathway level. The FT bladder proteome was best reflected by the detrusor transcriptome, consistent with the detrusor forming the bulk of the bladder wall. The urinary proteome correlated most strongly with the detrusor transcriptome at the gene level, whereas at the pathway level, and in particular in the direction of regulation, the closest agreement was with the mucosa. This is biologically plausible, since the urothelium is in direct contact with urine and contributes proteins through secretion, extracellular vesicle release and cell shedding, while proteins from deeper layers may reach the urine as urothelial barrier function becomes compromised after SCI (39). Thus, urinary proteins likely integrate signals from multiple bladder compartments, and bulk tissue analysis, while capturing the dominant detrusor response, may under-represent mucosa-specific changes.

Importantly, the increased expression of *GREM1*, *LTBP2*, *MSN* and *ACTA1* in bladder tissue from pediatric patients with neurogenic bladder supports the translational relevance of candidates identified in rodent models. Neurogenic bladder in this population is predominantly congenital, and tissue is typically obtained at reconstructive surgery and therefore reflects established disease, whereas the rodent models capture acquired obstruction over defined time courses. Evaluation of these markers in urine from well-phenotyped patient cohorts followed longitudinally will be required to establish their clinical utility.

In addition to the demonstration that clinically relevant interventions such as relief of obstruction and antimuscarinic medication modulated a subset of candidate biomarkers in tissue in the context of obstruction, we also showed that multiple urinary proteins induced by SCI were decreased with inosine treatment. Inosine was shown by us in previous studies to be effective at preventing oxidative DNA damage and PARP activation in the bladder, as well as bladder dysfunction including neurogenic detrusor overactivity (29, 35). Among the proteins increased in urine following SCI, but attenuated with inosine treatment, were Igfbp3 and Igfbp5. Although not studied in the context of bladder outlet obstruction, urinary levels of Igfbp3 have been linked to structural changes within the kidney in the transition from acute kidney injury to chronic kidney disease (40). In addition, increased expression of Igfbp3 has been implicated in progression of urethral fibrosis and stricture formation (41), highlighting a role for Igfbp3 in driving pathological alterations in tissue following injury. Igfbp5 has also been implicated in fibrosis in pre-clinical mouse models of kidney disease (42). In that study, endothelial Igfbp5, acting through the TGFβ1/Smad pathway, was shown to drive profibrotic changes in tubular cells in vitro, whereas genetic deletion of Igfbp5 either globally or in endothelial cells in vivo prevented these alterations. Our demonstration that inosine attenuated both Igfbp3 and Igfbp5 in the context of functional obstruction following SCI emphasizes the benefit of inosine in neurogenic bladder. In addition, these findings highlight the ability to capture meaningful changes in urinary proteins that reflect tissue-level alterations, including response to treatment.

Although the etiology of anatomic and functional obstruction is distinct, they share aspects of the pathophysiology including structural changes characterized by urothelial and smooth muscle cell proliferation and hypertrophy, oxidative stress and inflammation, fibroblast activation, extracellular matrix deposition, and tissue remodeling, all of which can contribute to functional decline (3, 4, 43). Among the biological processes and signaling pathways shown to be shared and enriched across different models of bladder outlet obstruction, processes associated with structural changes were prominent and included ECM-receptor interaction, cell/focal adhesion, regulation of the cytoskeleton, and cardiomyopathy reflecting conserved alterations to hollow organ remodeling across disparate insults. Consistent with pathway enrichment, of the 19 targets responsive to obstruction and treatment of obstruction, 8 are linked to the cytoskeleton and/or cytoskeletal remodeling.

The markers that showed consistent alterations in response to obstruction in multiple analyses --Cthrc1, Msn, Acta1, Ltbp2 and Grem1 --have all been linked mechanistically to cellular and tissue remodeling in a variety of contexts. Cthrc1 (collagen triple helix repeat containing 1) is a secreted glycoprotein that has been implicated in extracellular matrix turnover, fibroblast signaling and inflammation through modulation of TGF-beta and Wnt signaling (reviewed in (44)). Zhu and colleagues demonstrated a role for Cthrc1 in bladder remodeling following denervation injury in the rat (27). In that study, Cthrc1 was largely undetectable in healthy bladder tissue but increased markedly in response to pelvic ganglia cryoinjury.

Interestingly, in the same study the impact of bladder outlet obstruction on Cthrc1 levels was much more modest, highlighting differential sensitivity of Cthrc1 expression based on the nature of the injury. In our analysis, Cthrc1 was increased in rat bladder tissue in response to SCI. Furthermore, re-analysis of data from models in which obstruction was relieved surgically or with the anticholinergic agent tolterodine (26, 30) revealed that Cthrc1 expression was attenuated with treatment, consistent with a contribution to fibroproliferative remodeling in response to both anatomic and functional obstruction.

Moesin (Msn) is a component of the Ezrin-Radixin-Moesin complex that mediates interactions between the plasma membrane and actin cytoskeleton, and has been implicated in fibrosis in lung, liver and kidney (45–48). Studies in knockout mice have revealed conflicting results regarding the contribution of Msn to fibrosis in different organs. Exposure of Msnrrdeficient mice to bleomycin was associated with diminished repair and enhanced fibrosis compared to intact controls (45), consistent with a protective effect. In contrast, in models of acute liver injury, collagen deposition and migration of hepatic stellate cells were reduced in Msn-deficient versus-intact animals (46, 47). Furthermore, in a rodent model of unilateral ureteral obstruction (UUO), Msn levels were found to increase in the affected kidney tissue (48, 49) and urine (49) in parallel with the development of tubular injury and renal fibrosis. Chen and colleagues also showed that knockdown of Msn in renal tubule epithelial cells attenuated biochemical changes induced by the profibrotic factor TGFβ1 (48), suggesting that Msn promotes fibrosis. Consistent with these findings, we showed that Msn levels in tissue were associated with outlet obstruction of both neurogenic and non-neurogenic origin, urinary MSN was increased in response to SCI and both were sensitive to interventions associated with improved function.

Acta1 (skeletal muscle alpha-actin) is one of six actin isoforms and is predominantly expressed in skeletal muscle. However, its expression has been identified in the urinary tract during development (50), in activated fibroblasts (51) and in a variety of cancers, including those of epithelial origin (reviewed in (52)). Notably, re-expression of skeletal alpha-actin is a well-established component of the fetal gene program activated in pressure-overload hypertrophy of the heart (53) that, like the obstructed bladder, undergoes hypertrophic remodeling in response to increased outlet resistance. In the present study, increased *Acta1* expression was confirmed by qPCR in rat bladder after SCI and in human neurogenic bladder tissue relative to controls, and ACTA1 was detected among the differentially abundant proteins in the bladder mucosa of rats with pBOO and in the urine of SCI rats. Because actin isoforms share very high sequence identity, isoform-specific primers and proteotypic peptides are essential to attribute these signals specifically to ACTA1. Among the pathways identified in the models of obstruction evaluated in our study, ‘ECM-receptor interaction’, ‘Cytoskeleton in muscle cells’, ‘Cell adhesion molecules’ and ‘Focal adhesion’ were robustly enriched in all models. Consistent with their role as primary sensors of mechanical signals, integrins were found to be highly represented in multiple such pathways. Prior studies showed that integrin-mediated activation of RhoA increases SRF-dependent transcription of ACTA1 (54, 55), thereby providing a plausible mechanism for its identification as a candidate marker in our analyses.

Ltbp2 is one of a family of four latent TGFbeta-binding proteins, that play roles in production of extracellular matrix through regulation of TGFβ, organization of microfibrils and assembly of elastic fibers (reviewed in (56)). Expression of Ltbp2 was noted to be high in tissues enriched in ECM and exposed to mechanical stress, although the bladder was not examined in that study (57). Altered expression of Ltbp2 has been linked to fibrosis in a variety of organs including heart, lung and liver. LTBP2 was shown to be increased in cardiac tissue and plasma in patients with right ventricular dysfunction secondary to pulmonary arterial hypertension, characterized by progressive cardiac remodeling and functional decline (58). In that study, in addition to integrated transcriptomics and proteomics of cardiac tissue and plasma, the authors showed a similar increase in Ltbp2 protein in rat models of PAH compared to controls. In a separate study, Pang and colleagues showed that knockdown of Ltbp2 in a rodent model of dilated cardiomyopathy reversed multiple pathological endpoints including oxidative stress, myocardial fibrosis and cardiac dysfunction, in support of a causal role for Ltbp2 in driving cardiac fibrosis (59). These observations are analogous to the increased levels of Ltbp2 observed in rodent models of increased bladder outlet resistance following obstruction.

Grem1 has roles in fibroblast signaling, ECM turnover, and inflammation (reviewed in (60)). In addition to roles in development and cancer, Grem1 has also been implicated in fibrosis in different organs, including kidney, liver and lung, and specifically as a tissue-based biomarker in idiopathic pulmonary fibrosis (61–63) and inflammatory bowel disease (64, 65). Grem1 has been proposed to act through multiple mechanisms, including polarization of macrophages to an M2 phenotype and by promoting epithelial-to-mesenchymal transition with ensuing ECM deposition (66, 67). Although not explored in remodeling associated with bladder outlet obstruction, Grem1 is expressed in the bladder where it has been proposed as a contributor to and biomarker for bladder cancer progression and prognosis (68, 69). Interestingly, in those studies, gene ontology and pathway analyses of bladder cancer tissues linked GREM1 to structural changes including collagen production and collagen-ECM interaction, consistent with both our observations across different models of obstruction and with a role in tissue remodeling.

CTHRC1, MSN and GREM1 have been identified previously in urine but their potential as urinary biomarkers of outlet obstruction has not been explored. Urinary CTHRC1 levels were explored in patients with myasthenia gravis, an autoimmune disorder characterized by disrupted neuromuscular communication (70). In that study, CTHRC1 levels declined in the urine of individuals with myasthenia gravis compared to healthy controls, although a mechanistic role was not investigated. Increased MSN levels were identified by urine proteomic analysis in response to tubular injury following UUO (49), whereas GREM1 was found to be elevated in urine in a subset of patients with renal damage associated with glomerular disease (71). These latter findings raise the possibility that MSN and GREM1 detected in our analyses are of renal origin. However, our multiomics analysis, together with validation in bladder tissue from rodents with functional obstruction is consistent with urinary proteins originating, at least in part, from bladder tissue and reflecting its state. Nevertheless, in neurogenic bladder, high intravesical pressure and vesicoureteral reflux can cause upper tract damage, so a renal contribution cannot be excluded. These observations emphasize the need in future studies for orthogonal validation to ensure sensitive, specific detection of putative disease biomarkers.

There are several limitations of this study. First, bladder function was not assessed in the studies from which transcriptomic data were obtained, therefore the relationship between candidate biomarkers and functional status cannot be determined. Second, a variety of rat strains, and animals of both sexes were used to generate models of outlet obstruction. As a result, we cannot appreciate sex differences in differential gene/protein expression associated with obstruction and/or remodeling. Third, the timeframe over which bladder pathology develops in experimental models of acute obstruction versus that occurring in humans is substantially different, highlighting the need for validation in well-phenotyped human cohorts in future studies. Fourth, our own proteomic data, including the urine proteome, were derived from a single model of neurogenic obstruction; although two public proteomic datasets from rats with pBOO detected a subset of candidates at the protein level, these comprised few biological replicates, and the correlations between tissue transcriptome and urine proteome observed after SCI may not apply to other forms of obstruction. Fifth, the number of animals per group in our SCI omics analyses was small (n = 3-4), limiting statistical power to detect modest changes. Finally, several candidates, including Grem1, Cthrc1 and Igfbp5, are also altered in bladder cancer and in fibro-inflammatory conditions; their specificity for obstruction will need to be established in appropriate disease-control cohorts, ideally as part of a multi-marker panel.

In summary, we have identified a panel of biomarkers from rodent models that reflect the response to bladder outlet obstruction over time, and that are sensitive to surgical or pharmacological treatment. Our results provide a platform for prospective evaluation of candidate biomarkers in humans with bladder outlet obstruction.

## Materials and Methods

### Sex as a biological variable

The animal model generated in our laboratory previously (29) was performed in male rats to enable comparison with prior studies in our laboratory. Publicly available data from animal models was from male and female rats. For human tissue analysis, specimens from both male and female subjects with neurogenic bladder and controls were assessed. Sex as a biological variable was not considered in this study.

### In silico analyses of transcriptomic data

Publicly available datasets from rodent models of bladder outlet obstruction were re-analyzed to identify transcriptomic changes in response to injury. Data were retrieved from the NCBI GEO Database using GEOQuery v2.72.0 (72). Six datasets were retrieved: GSE47080 (26), GSE104540 (27), GSE197544 (28) GSE294927, GSE294930 (29) and GSE128618. Data quality was assessed using PCA plots to determine how well data from biological replicates for a given mode of injury clustered together using the base R stats package (73), and therefore supported differential gene expression analysis **(Suppl. Fig. 1A-H)**. Microarray data (GSE104540 and GSE47080) were analyzed using linear models for microarray analysis (LIMMA) package v3.60.6 in R (74). For RNAseq data, SRA raw files were downloaded from the SRA run selector using the SRAtoolKit v2.10.7 (75), genome indexes were generated using the most recent versions of the rat primary assembly, Rnor_6.0 (for consistency with our previous analysis) (29), and aligned to the rat library using STAR aligner (76) v2.7.9a, using the following settings; i) GeneCounts quant mode, ii) Generated BAM files sorted by coordinate, iii) and paired or unpaired reads depending on the fastq files retrieved from the SRA. Count matrices were generated using the Subreads, FeatureCount function v2.0.3 (77) without strand specificity. The resulting count matrices were analyzed with DESeq2 (78) v1.44.0, using the following parameters to consider genes for downstream analysis: i) likelihood ratio test; ii) FDR < 0.05; iii) genes with an absolute value of log2FC >0.5. Data frame manipulations were performed using a variety of packages in R including tidyR v1.3.2 (79), dplyR v1.1.4 (80), tibble v3.3.1 (81), and stringR v1.6.0 (82). Plots were constructed using ggplot2 v4.0.1 (83) and GGally v2.4.0 (84). Conversion of Ensembl and Affymetrix microarray gene IDs was performed using a combination of biomaRt v2.60.1 (85) and org.Rn.eg.db v3.19.1 (86). Venn diagrams of overlapping genes were generated with ggVennDiagram v1.5.7 (87) and Volcano plots were generated using EnhancedVolcano v1.22.0 (88). Pattern analysis of DEGs was performed using the DEGreport v1.40.1 package (89). ClusterProfileR v4.12.6 (90) and pathfindR v2.3.1 (91) were used to perform enrichment analysis using GO terms for molecular function, biological process and cellular compartment on DEG lists without and with Log2FC and adjusted p-values, respectively. Hierarchical clustering of enriched terms was performed to cluster similar groups of terms together. Heatmaps were generated using pheatmap package v1.0.13 (92).

### Integrated multi-omics and correlation analyses

Differentially expressed genes (DEGs) and proteins (DEPs) were matched by gene symbol. For each pairwise comparison between datasets and time points, Pearson correlation of log2 fold changes was calculated using (targets differentially expressed in both datasets of the pair / the 161 targets differentially expressed in all datasets at all time points), and the coefficient of determination r and P value are reported. KEGG pathway enrichment was performed separately for each list of DEGs or DEPs, and fold enrichment scores of pathways enriched in both datasets were compared by correlation analysis. To capture the direction of regulation, a relative regulation score was calculated for each enriched pathway as (number of up-regulated targets-number of down-regulated targets) / (total number of differentially expressed targets in the pathway) x 100, and scores were compared between datasets by correlation analysis.

### In silico analyses of proteomic data

Two publicly available proteomic datasets were retrieved from the PRIDE Archive, PXD007571 and PXD029337. These were processed separately as the platforms used to generate them and the format they were submitted to PRIDE in differed. The PXD007571 dataset comprised.mzid and.mgf files, which were downloaded and processed into data frames and Spectra objects with the Spectra v1.22.2 (93) respectively. Spectrum identifiers were extracted from the Spectra object and used to filter and group the PSMs from the mzid files into protein groups for each sample. A proxy for protein abundance was generated by counting the MS2 spectra assigned to each protein group based on the fragment events in thes.mgf spectral file. The number of distinct PSMs and the total number of PSMs for each protein group were recorded. Quantified protein groups were kept for downstream analysis if they were present in >=50% of the 4 samples. The summed PSM counts were converted to a log2 transformed abundance matrix prior to downstream analysis.

The second proteomic dataset comprising full thickness bladder tissues following 2 weeks of pBOO or sham controls (PXD029337) was limited to raw.wiff and.wiff.scan files. These proprietary formats were converted to the universal.mzML format using MSConvert [PMID: 28188540] with the peak picking filter based on the vendor algorithm. The converted.mzML files were then subjected to peptide-based quantification using Fragpipe under a modified protocol to address iTRAQ-8plex methodology used to generate the data. The pooled sample data was extracted from each of the 15 files using the relative isotopic labels 113-119 and 121. Quality control analysis informed the removal of samples labeled with 113, 114 and 115 isotopic tags. Further, supplemental data from the original publication informed the identity of 3 isotopic labels (119, 117 and 118). Additional quality control assessments informed the identity of samples labeled with 121 and 116. The absence of complete metadata for these data limits the certainty of the resulting data. Following demultiplexing, peptides not unique to a single protein were removed. The resulting unique peptides were log2 transformed prior to downstream analyses. Both PXD007571 and PXD029337 datasets were limited to an average of two biological replicates across each of two conditions. This required the use of LIMMA based tools including the empirical Bayes moderated t-test to assess significance using a model built using all samples.

### Preparation of tissue and urine samples from rat SCI model

Full-thickness bladder tissues from rats without or with SCI for 2, 8 or 16 weeks were generated as described previously (29) were lysed in 50mM NH_4_HCO_3_/2% SDS using FastPrep Lysing matrix S beads (MP Biomedical) and quantified by NanoDrop. Tissue lysates and urine samples from the same animals were centrifuged at 4000 rpm (680 x g) for 10 minutes to remove cellular debris. Samples were concentrated in 10kDa cut-off centrifuge filters and washed twice with 8M urea in 100mM triethylammonium bicarbonate (TEAB). Proteins retained on the filter were reduced in 10 mM DTT, alkylated in 25mM iodoacetamide, and then subjected to digestion with trypsin in 50mM NH_4_HCO_3_. Resulting peptides were collected by centrifugation, washed with 20% acetonitrile (ACN), recentrifuged and washed prior to final centrifugation with 70% ACN. Peptides were loaded onto a third-generation Aurora Ultimate reverse-phase nLC column (IonOpticks) using an Easy-nLC 1200 (Thermo) and analyzed via DIA-MS using an Orbitrap Exploris 240 MS (Thermo). Detailed LC-MS/MS parameters are presented in supplementary materials.

### qRT-PCR validation

Expression of a subset of genes identified as differentially expressed across all datasets was validated by qRT-PCR. For validation, primers were designed for rat and human transcripts using NCBI primer design tool such that the length of the product was between 100-300 bases to facilitate rapid detection via qPCR. RNA was isolated from tissues generated in-house essentially as described. Briefly tissues were homogenized in 500µL of TRIzol. 100µL of chloroform was added to the TRIzol, mixed by vortexing and centrifuged for 15 mins at 7826 x g. The aqueous phase was added to an equal volume of 70% ethanol before using the RNeasy minikit (Qiagen) to isolate RNA following the manufacturer’s protocol. RNA quality was inferred using NanoDrop based on the 260/280 ratio. Samples were discarded if the ratio was beyond the 1.90-2.1 range. cDNA was generated using the iScript cDNA synthesis kit (Bio-Rad) following the manufacturer’s protocol and amplified using gene-specific primers (**Suppl. Table 1**). Primers were generated with the NCBI primer design tool using the following criteria; i) primers must span exons, ii) melting temperature = 60°C +/-2°C, iii) GC content = 50% +/-3%, iv) amplicon size 100-500bp, v) unintended amplicons limited to those with >= 98% homology or limited, when possible, to predicted and known isoforms. Primers, cDNA and SYBR Select master mix (ThermoFisher) were combined in a 96-well PCR plate and run in a QuantStudio3 thermocycler (Thermo Fisher). Expression of each target was normalized to the housekeeping genes Gapdh and Rps18. Several housekeeping genes including Sdha, Abcf1, Gapdh, Rps18, and Actb were analyzed initially, with Gapdh and Rps18 emerging as the least variable, and therefore selected for normalization. Amplification of a single product was verified by melt-curve analysis at the conclusion of each run.

### Immunoblot analysis

Protein samples prepared from rodent urine were generated following acetone precipitation, desiccation and resuspension in cell lysis buffer (20 mM Tris (pH 7.5), 150 mM NaCl, 1mM EDTA, 1 mM EGTA, 2.5 mM NaPPi, 1 mM β-glycerophosphate, 1 mM Na_3_VO_4_, 1µg/ml leupeptin). Protein concentrations for rat urine protein samples were quantified using the MicroBCA assay (ThermoFisher). Fifty micrograms of rat urine protein samples were resolved by SDS-PAGE, electrotransferred to nitrocellulose membranes with protein transfer verified by Ponceau S staining. After incubation in 10% non-fat dried milk in PBS/0.1% Tween-20 (PBS-T) for 1 hr to block non-specific binding sites, membranes were washed in PBS-T, and incubated in primary antibody (GREM1 (#sc-515877, Santa Cruz), MSN (#3146S, Cell Signaling Technology), CTHRC1 (#sc-293270, Santa Cruz) or LTBP2 (#sc-166199, Santa Cruz))(**Suppl. Table 2**) overnight at 4°C. Membranes were washed 3 times for 15 mins in PBS-T prior to incubation in species-specific HRP-conjugated secondary antibody for 1 hr at room temperature in 10% non-fat powdered milk/PBS-T. Signals were visualized by incubation of membranes in SuperSignal™ West Pico PLUS Chemiluminescent Substrate (Thermo Fisher) following the manufacturer’s protocol and capture by ChemiDoc chemiluminescence imaging or on HyBlot CL™ Autoradiography Film (Thomas Scientific). Films were digitized using an Epson scanner and quantified with FIJI.

### Statistical analysis

For qPCR and immunoblot analysis, 3 biological replicates were assessed at 3 time points for both control and SCI conditions in rat. 8 biological replicates (n=4 male, n=4 female) were assessed for each of the detrusor and mucosa sub-compartments of bladder tissues from pediatric patients with neurogenic bladder and controls. Data were assessed for normality and paired-Wilcoxon signed-rank test was used to calculate p-values. * p-value < 0.05, ** p-value < 0.01, *** p-value < 0.001. Statistical significance was assessed in microarray and proteomic data using empirical Bayes moderated t-tests implemented in LIMMA (94, 95), with multiple testing correction performed using the Benjamini-Hochberg method (96)(* adjusted p-value < 0.05).

### Study Approval

The animal experiments conducted in this study were performed in strict accordance with the recommendations provided in the Guide for the Care and Use of Laboratory Animals of the National Institutes of Health. The experiments were approved by the Animal Care and Use Committee at Boston Children’s Hospital (protocol # 16-08-3256R). Bladder tissue was obtained from male and female pediatric patients with neurogenic bladder (n = 4 in each group; age (0-300 months)), at the time of bladder augmentation and from age-and sex-matched control patients without bladder dysfunction. Detrusor and mucosa were separated by microdissection and stored at-80C. Tissues were obtained under an IRB-approved protocol at Boston Children’s Hospital (X06-05-0271), with written informed consent received prior to participation.

## Data availability

Data generated by our lab are available in public databases. The mass spectrometry proteomics data from rat tissue and urine have been deposited to MassIVE, a contributor to the ProteomeXchange Consortium, with the dataset identifiers MSV000101848 (PXD046096, PXD078368). RNA-seq data are available in NCBI GEO under accession numbers GSE294927 and GSE294930. Values for all data points in graphs are provided in the file Supporting Data Values File.xlsx that accompanies the article.

## Author contributions

A.B-A curated the data, performed bioinformatics analysis with supervision by A.H.G., sample preparation with assistance from K.C. and S.D., and immunoblotting and qRT-PCR assays with assistance from G.L.O and J.K. A.B-A analyzed data, generated figures and wrote the original draft of the manuscript with R.M.A; mass spectrometry data were generated by Y.T. and J.W.F, with data analysis by B.D., Y.T. and J.W.F. R. Lee supervised the study and R. Adam conceived and supervised the study, and wrote the manuscript.

## Supporting information

Supporting Data Values

## Acknowledgements

The study was supported by National Institutes of Health grants R01 DK077195 (RMA) and R01 DK127673 (RSL, RMA) and by the Children’s Urological Foundation.

This work was supported by grants from the National Institutes of Health (R01 DK077195 (RMA), R01 DK127673 (RSL, RMA)), and the Children’s Urological Foundation.

