## Supplemental Figures and Tables for "Integrated transcriptomic and proteomic analyses identify novel biomarkers of bladder outlet obstruction"

Alexander A. Bigger-Allen^1,2^, Barnali Das^1,2^, Yang Tang^1^, Kyle Costa^1^, Gabriel Luis Ocampo^1^,

Ali Hashemi Gheinani^3,4^, Shannon DiMartino^1^, Jane Kaull^1^, John W. Froehlich^1,2^, Richard Lee^1,2,*^, Rosalyn M. Adam^1,2,*^

^1^Urological Diseases Research Center, Boston Children’s Hospital, Boston, MA, USA.

^2^Department of Surgery, Harvard Medical School, Boston, MA, USA.

^3^Functional Urology Research Group, Department for BioMedical Research DBMR, University of Bern, Switzerland.

^4^Department of Urology, Inselspital University Hospital, 3010 Bern, Switzerland.

**Supplementary Materials**

Supplementary Methods

Supplementary Figures 1-9 and Legends

Supplementary Tables 1 and 2

**Supplementary methods**

**LC-MS/MS analysis**

LC-MS/MS analysis was performed on an EASY-nLC 1,200 coupled to an orbitrap Exploris 240 mass spectrometer via a Nanospray Flex ion source (all Thermo Fisher Scientific). Purified peptides were separated at 60 °C using a 25 cm × 75 μm (i.d.) third-generation Aurora Ultimate reverse-phase nanoLC column (IonOpticks). Mobile phase A consisted of water with 0.1% formic acid (v/v) and mobile phase B consisted of 80% acetonitrile, 20% water, and 0.1% formic acid (v/v/v). The flow rate was maintained at 400 nL/min throughout the separation. Solvent B started at 2% and was increased linearly to 35% over 50 min, then further increased to 40% B over 10 min, followed by a rapid ramp to 95% B in 2 min, with a subsequent 2 min isocratic wash at 95% B at the end. MS data were acquired in the data-independent acquisition (DIA) scan mode for rat tissue samples, including four biological replicates per sample type/condition. Full-scan MS1 spectra were acquired in the m/z range of 400–1,000 at a resolution of 90,000 (specified at *m/z* 200), with the automatic gain control (AGC) set to a target value of 3 × 10⁶. Each full MS scan was followed by 24 static MS/MS scans with an isolation window of *m/z* 25 and an overlap of *m/z* 1.0, covering the same *m/z* range (400–1,000) at a resolution of 22,500 (at m/z 200). Precursor was fragmented by higher-energy collisional dissociation (HCD) with a normalized collision energy of 30%. Fragment ions were accumulated until reaching an AGC target of 5 × 10^5^ or a maximum accumulation time of 46 ms.

**Mass spectrometry data processing**

MS DIA data were processed in Spectronaut using directDIA mode with default settings. Searches were performed against the Swiss-Prot rat reference proteome (UniProt ID: UP000002494; release 2025-09-08; 47,914 entries). Carbamidomethylation on cysteine (C) was set as a fixed modification, while and acetylation of the protein N-terminus and oxidation of methionines (M) were selected as variable modifications.

**Supplementary Figures**

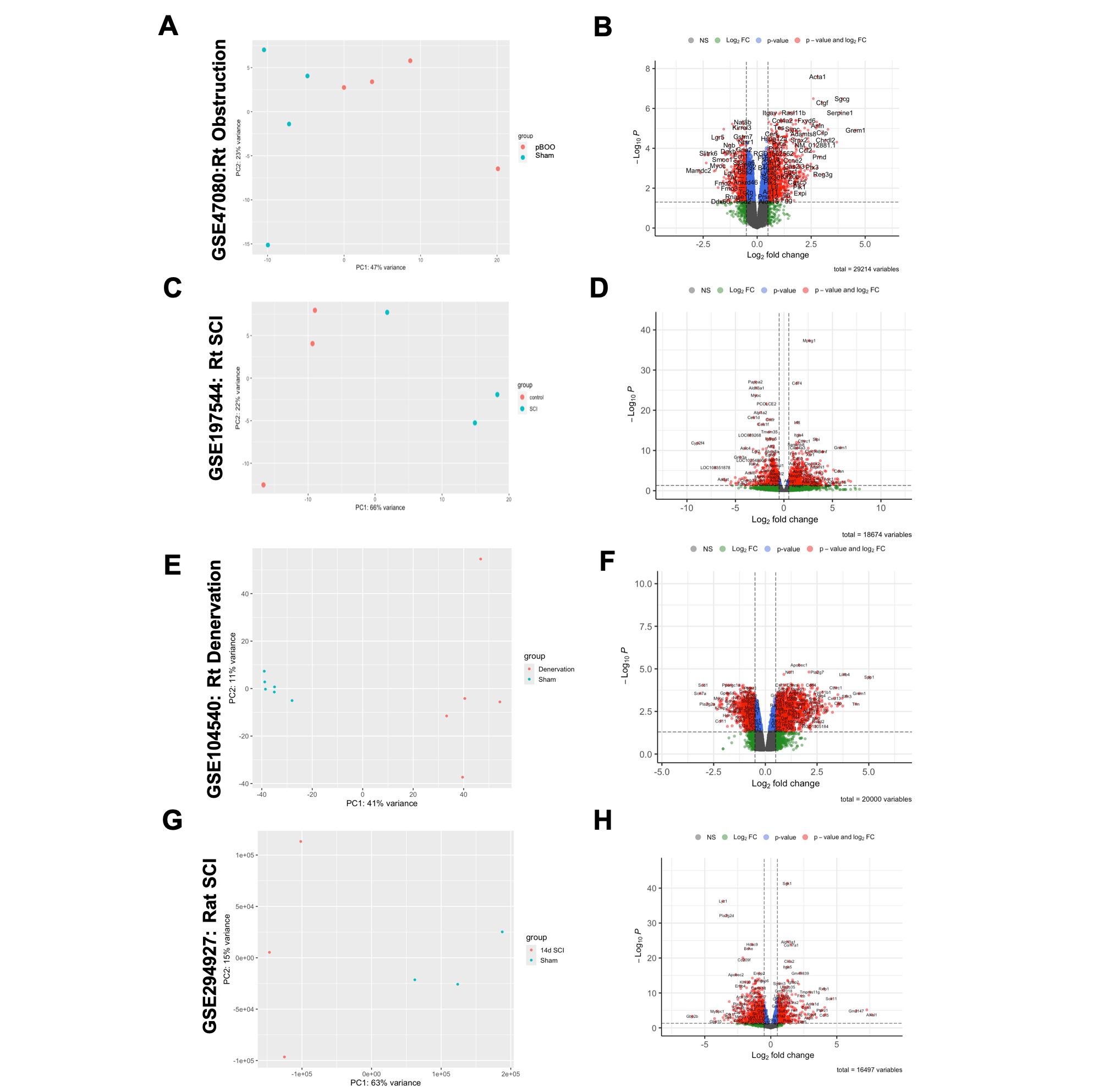

**Supplementary Figure 1. PCA and Differential expression analysis of rodent bladder injury datasets.**

**(A, C, E, G)** Principal component analysis of the raw expression data from each of 4 datasets representing bulk RNA from bladders following injury. The first two principal components are represented in each PCA plot and represent the greatest variation in each dataset. In four models of rat bladder obstruction, injury accounts for more than 40 percent of the variation in the dataset separating the injured samples from the control (PC1 on the x-axis).

**(B, D, F, H)** Volcano plots depicting the results of the differential expression analysis in each of four datasets. Each dot represents a single gene. Genes are color coded gray if they have a –log10(p.adj.) less than 1.3, and their absolute log2FC is less than 0.25. Genes are color coded green if they have a –log10(p.adj.) less than 1.3 and their absolute log2FC is greater than 0.25. Genes are color coded blue if they have a –log10(p.adj.) greater than 1.3 (p.adj < 0.05) and have an absolute log2FC < 0.25. Genes of interest for downstream analysis are labeled in red and have a –log10(p.adj) greater than 1.3 and have an absolute log2FC >0.25.

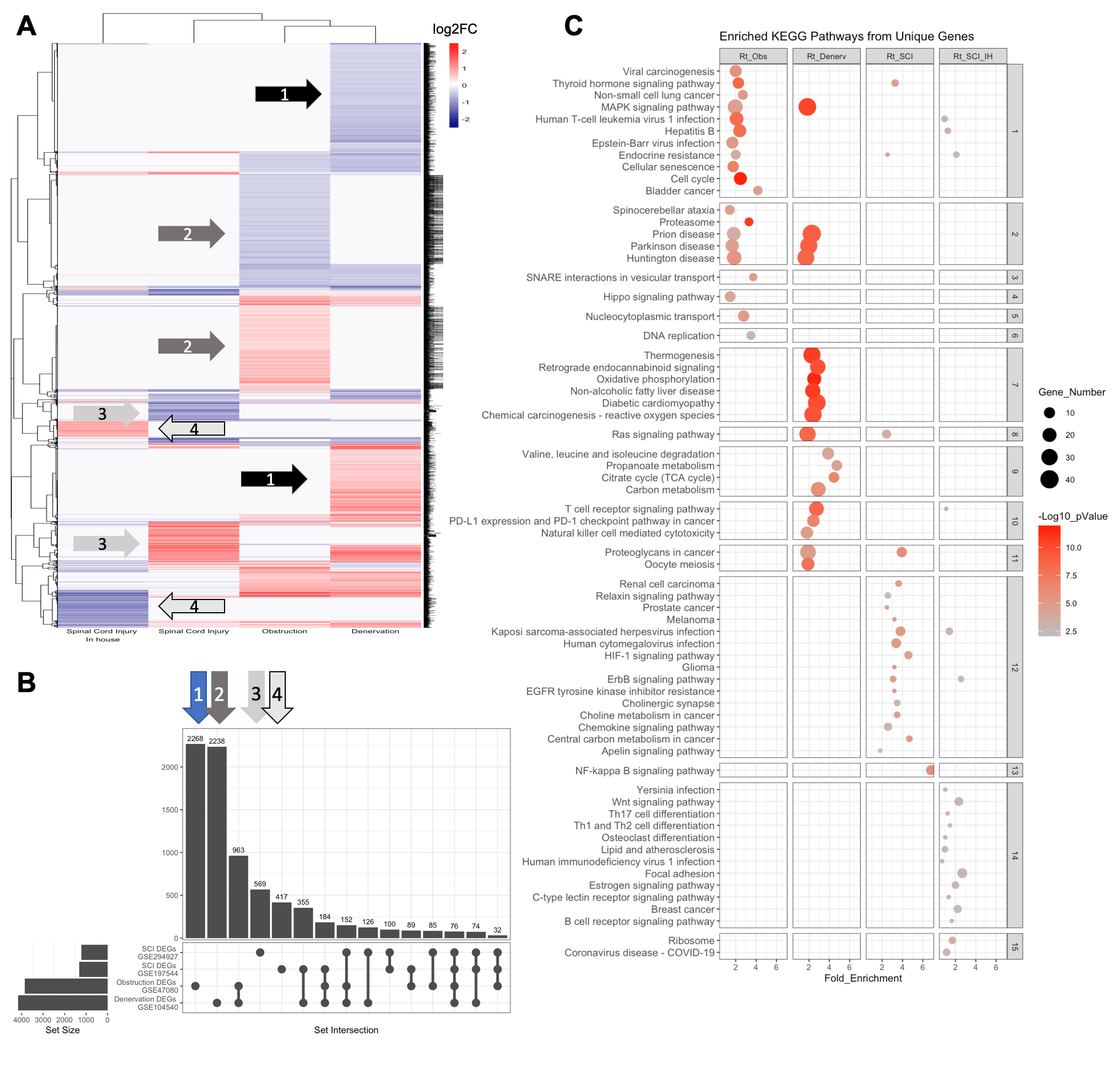

**Supplementary Figure 2. Unique DEGs and enriched pathways in rat models of bladder obstruction.**

(**A)** Heatmap of the log2FC of all differentially expressed genes from each of 4 datasets comprising rat models of partial bladder outlet obstruction, denervation, or spinal cord injury (SCI) compared to corresponding controls. Hierarchical clustering using Pearson correlation on the rows and columns (genes and models) demonstrates groups of DEGs that are unique to each of the four datasets (labeled arrows). **(B)** Upset plot of the groups of genes uniquely expressed in each data set highlighted in panel A shows the number of DEGs unique to or shared between the four datasets. **(C)** KEGG pathway enrichment analysis for unique DEGs from each dataset: bladder outlet obstruction (Rt_Obs), denervation (Rt_Denerv), or spinal cord injury (SCI).

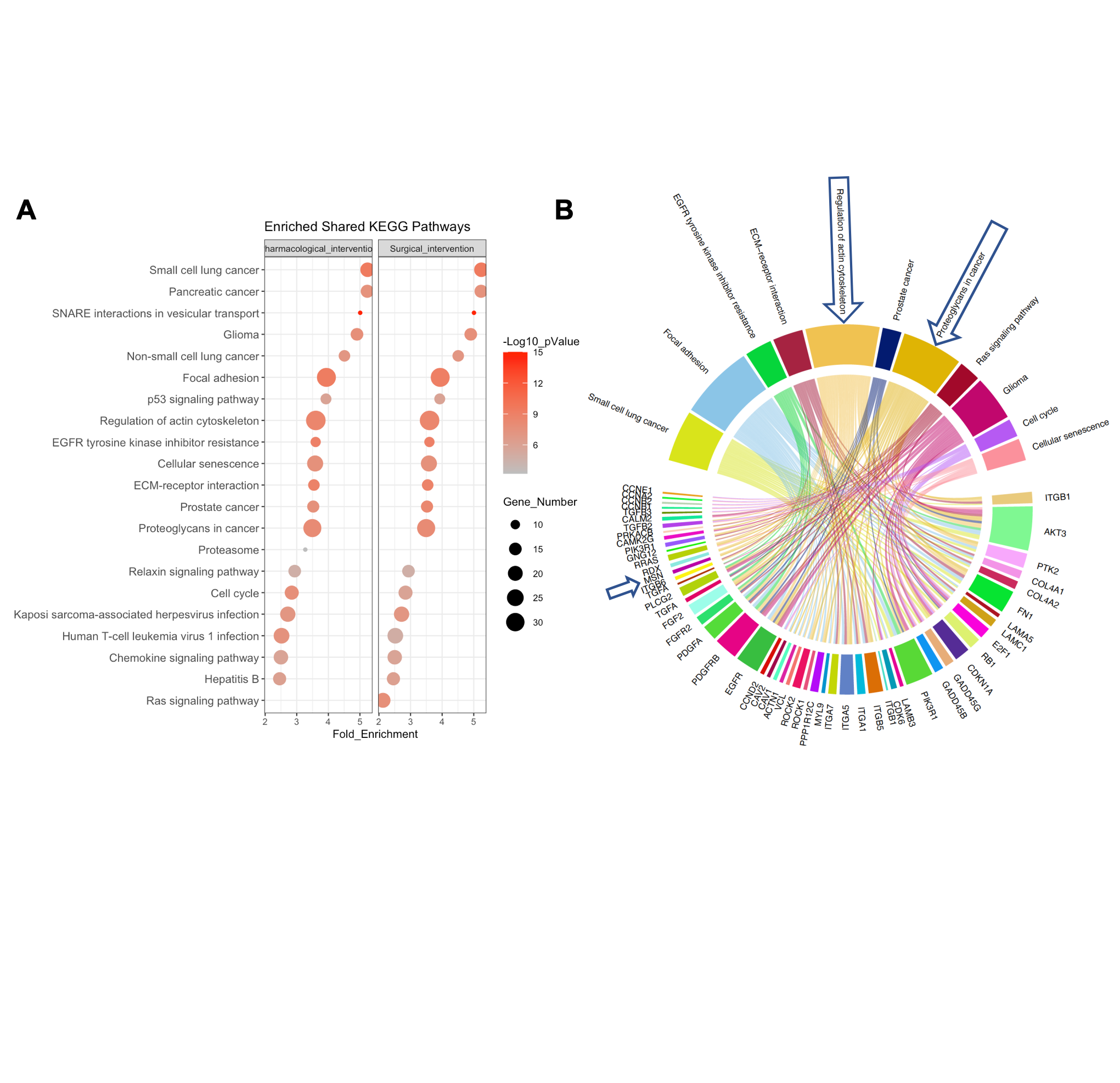

**Supplementary Figure 3. Pathway analysis and associated DEGs shared in pBOO datasets**

**(A)** KEGG enrichment analysis of shared DEGs between GSE47080 and GSE128618. **(B)** Chord diagram connecting the shared enriched pathways between both intervention datasets and the associated DEGs. Blue arrows highlight Msn and the KEGG pathways it is assoicated with.

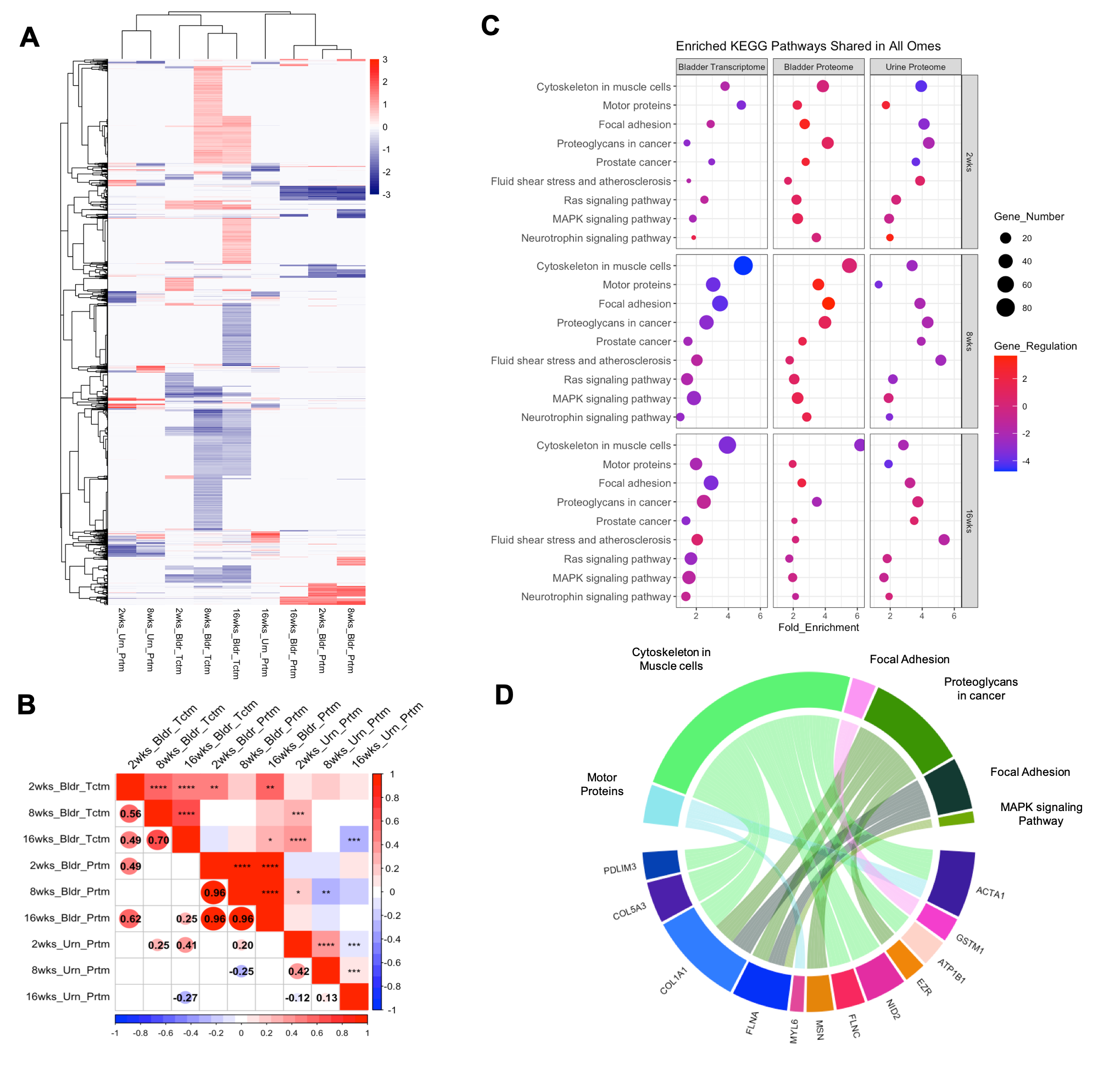

**Supplementary Figure 4. Multi-omic comparison in time course of SCI in the rat bladder and urine**

**A)** Heatmap of the log2FC of all 6840 unique targets across all “omes” at all time points. Hierarchal clustering of rows and columns using 1-pearson correlation was employed to quantify the similarities between each ome. **B)** Correlation plots quantify the extent to which each time point of each ome correlates. Top half of correlation plot indicates the extent of statistical significance in the correlation of each time point ome. The bottom half explicitly shows the R^2 value between the omes that have statistically significant correlation. **C)** Dotplot of all shared enriched KEGG pathways across all time-points and omes using the DEGs and DEPs of spinal cord injured animals compared to controls. Dot size reflects the number of genes in each pathway, Dot color reflects the ratio of up- and down-regulated targets in each dot. All terms were enriched with an FDR <= 0.05. **D)** A chord diagram connecting the targets shared in all omes in at least one time point and their associated KEGG pathways.

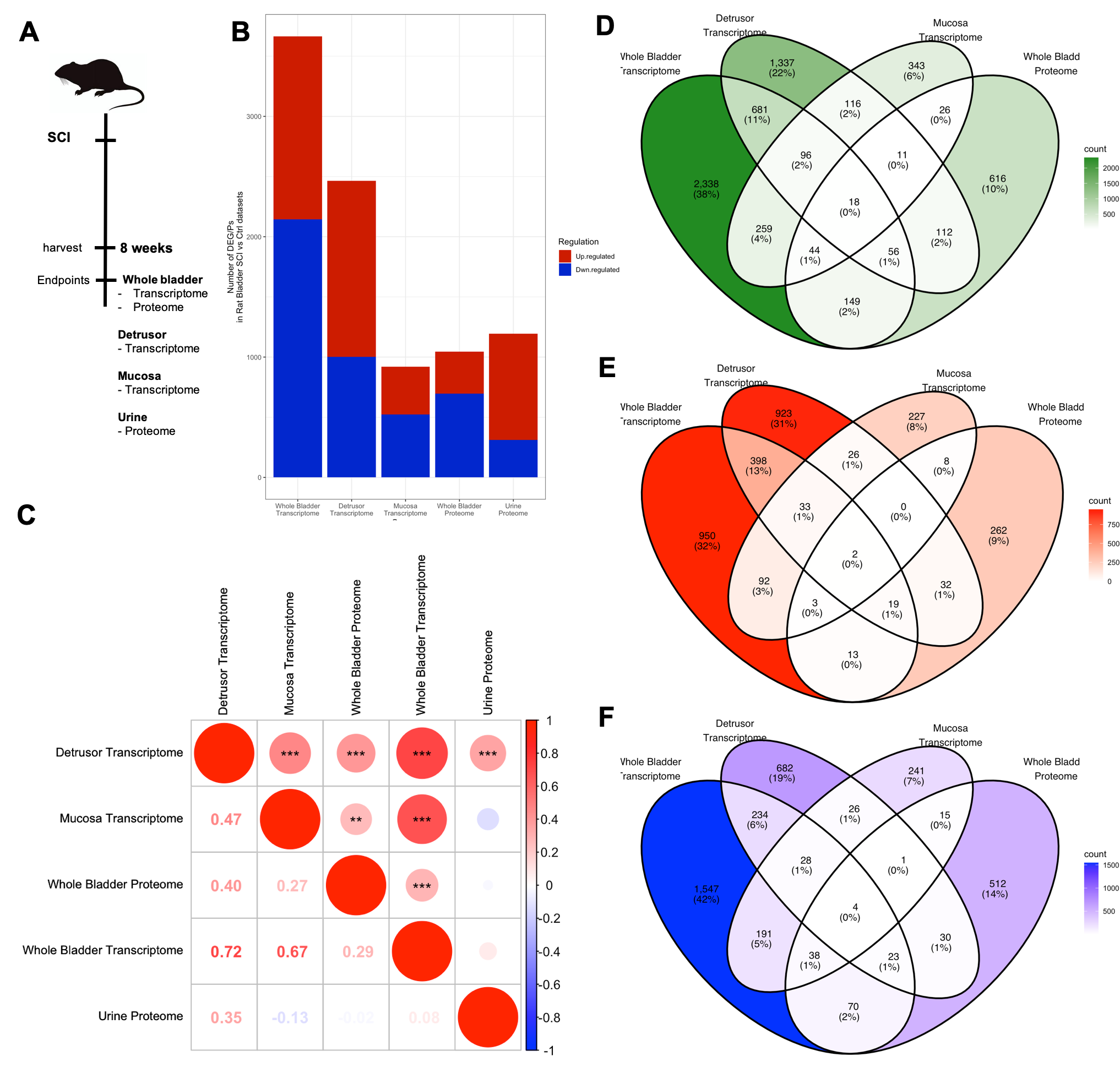

**Supplementary Figure 5. Correlation of transcriptomic and proteomic changes induced by SCI**

**A)** Proteomic and transcriptomic analyses were performed in whole bladder tissue, Detrusor and Mucosa separated tissue or urine of rats following 8 weeks of SCI. **B)** A bar chart compares the four sample types highlighting the variation in the total number of differentially expressed targets. **C)** Pairwise correlation analysis identified statistically significant correlation between the whole bladder and both the Detrusor and Mucosa transcriptome (r = 0.724 and r = 0.671, p<0.001). The whole bladder proteome showed a negative correlation with the detrusor transcriptome (r2 0.724, p<0.001). **C)** Venn diagram compares differentially expressed targets regardless of direction of expression. **D)** Venn diagram compares only the up-regulated targets differentially expressed in each analyte**. E)** Venn diagram compares only the down-regulated targets that are differentially expressed in each analyte

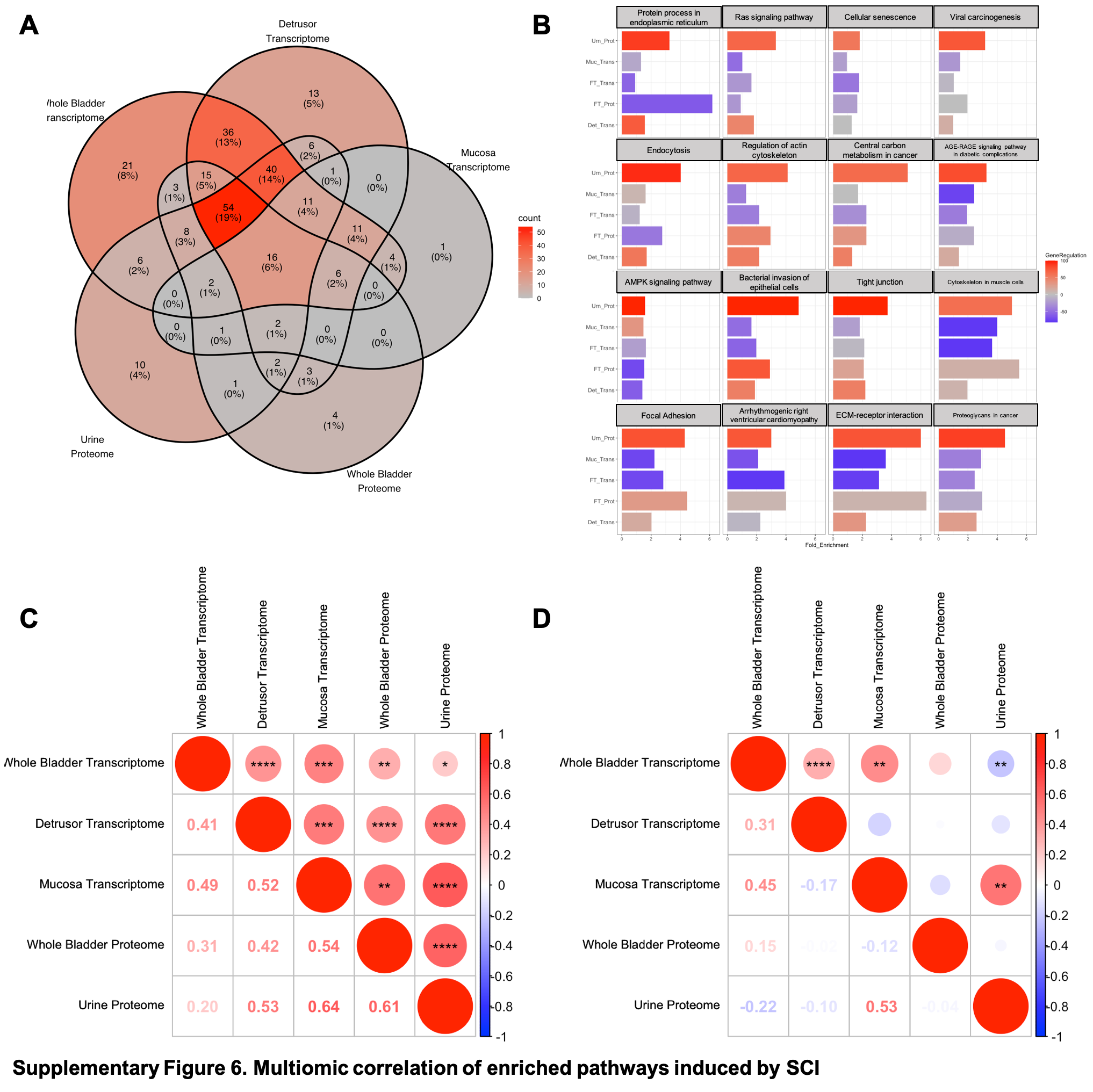

**Supplementary Figure 6. Multiomic correlation of enriched pathways induced by SCI**

**Enrichment analysis was performed on the differentially expressed targets in each ome and analyte.**

**A)** Venn Diagram highlights the extent to which the KEGG pathways enriched in each ome are shared. **B)** The column chart shows the 16 enriched KEGG pathways (from the center of the venn diagram in A) that are shared across all five DEG/P lists. Columns are color coded based on the percentage of up- and down-regulated targets associated with each kegg pathway for each ome-analyte. More up-regulated or down-regulated DEGs is color-coded as increasing red or blue, respectively. **C)** Pair-wise correlation analysis of all enriched kegg pathways between any two ome-analyte DEG/Ps list. Fold enrichment is used as the numeric value for this analysis. **D)** Pair-wise correlation analysis of all enriched kegg pathways between any two ome-analyte DEG/Ps list. Gene regulation, the realitve proportion of up- and down-regulated targets is used as the numeric value for this analysis.

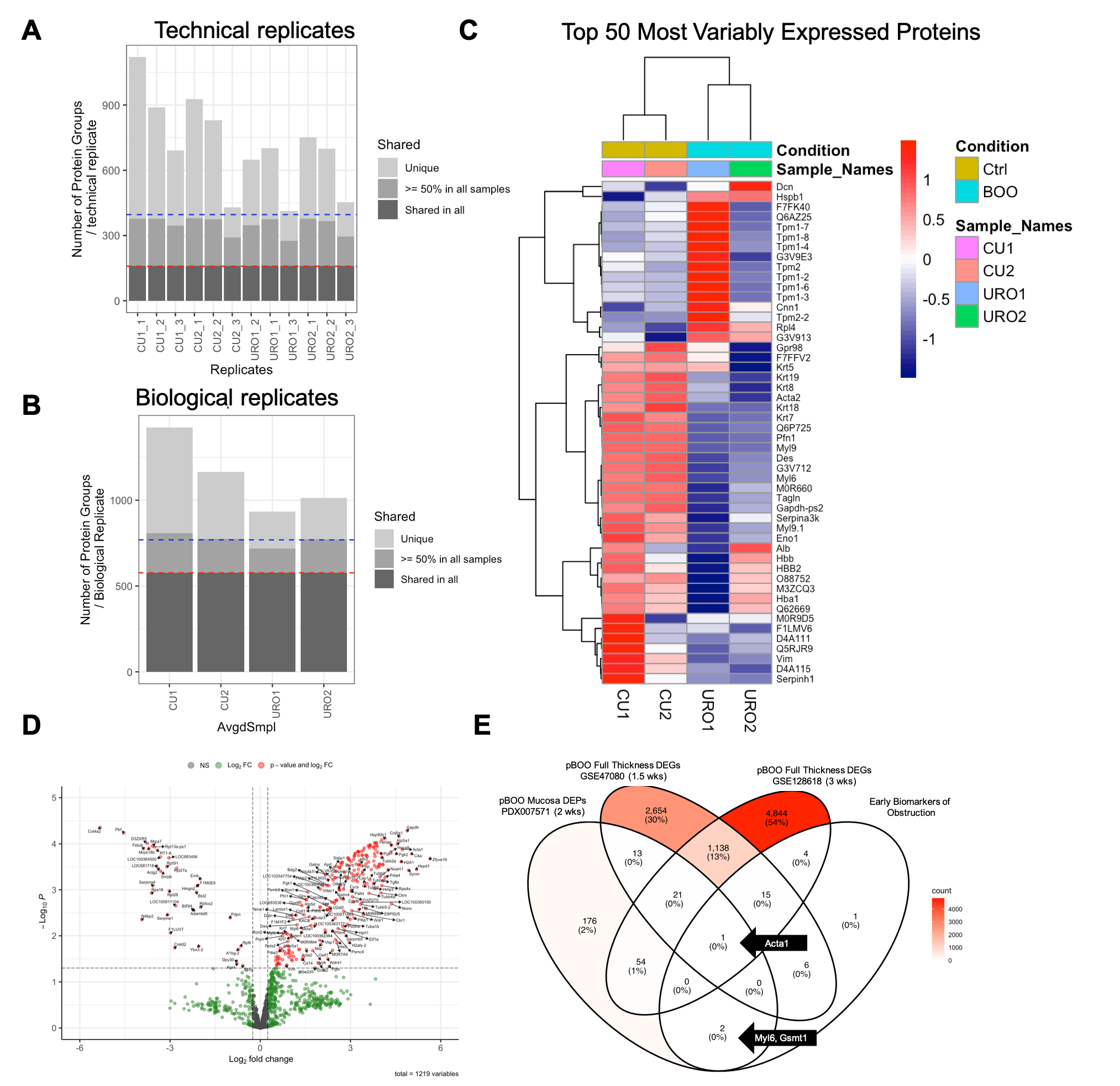

**Supplementary Figure 7. Identification of protein biomarkers in rat urothelium following pBOO**

Reanalysis of proteomic data from PRIDE database PXD007571 comprising bladder urothelium from control (CU) and pBOO (URO) rats. **A)** Bar chart highlighting variation in protein group detection among technical replicates and across conditions. 150 protein groups were detected in all runs and ~350 protein groups identified in 50% of the runs. **B)** Bar chart highlighting the results of the total number of protein groups shared in all (~510) or in >=50% (~700) of the biological samples after merging the protein groups for each technical replicate. **C)** A heatmap of the top 50 most variably expressed proteins across the biological replicates, highlighting intra- and inter-condition variation. Targets such as Dcn, Tpm1/2 and associated isofroms, Krt5, Alb, Hbb, Hbb2, and Hba1 show high variability between the pBOO replicates. Variability between control replicates is highlighted in Alb, and M0R9D5 (Ahnak). The color of each cell reflects a row-wise scaling of the spectral counts for each protein group. **D)** Volcanoplot shows roughly 200 DEPs that have a |log2FC| > 0.25 and adjusted p-value < 0.05 highlighted in red. Green dots represent proteins that met the log2FC cutoff but not the pvalue cutoff. Gray dots represent the proteins that did not meet any cutoff. **E)** VennDiagram of DEPs from rat mucosa proteome following 2wks of pBOO in overlap with 54 and 75 DEGs in full thickness rat bladder transcriptomes following 1.5 and 3 weeks of pBOO. 3 DEPs overlapped with the predicted early biomarkers of obstruction.

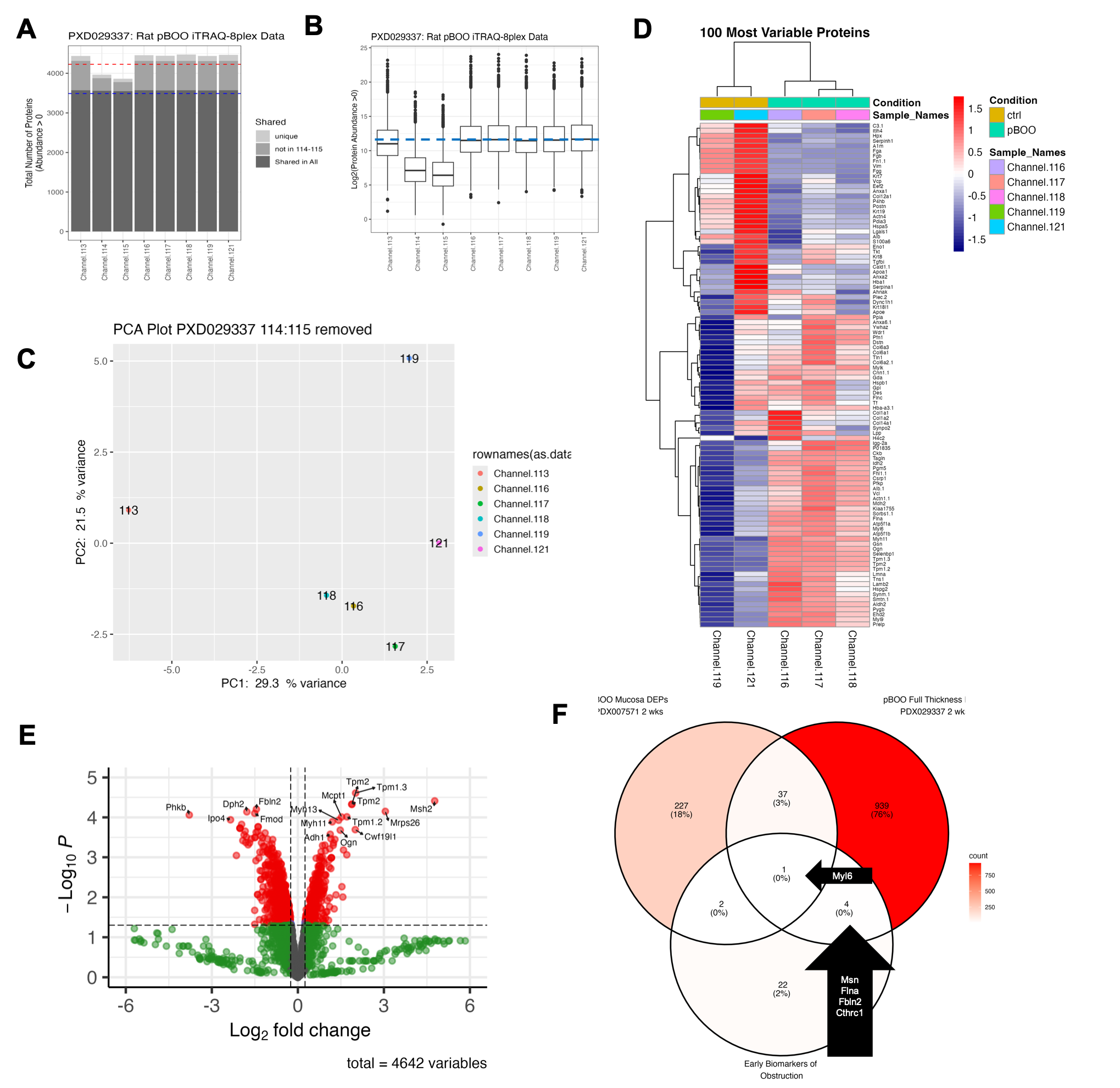

**Supplementary Figure 8. Identification of protein biomarkers in rat full thickness bladder following pBOO**

Reanalysis of proteomic data from PRIDE database PXD029337 comprising full thickness bladders from control and pBOO rats that were multiplexed with an iTRAQ-8plex labeling kit. **A)** Bar chart highlighting variation in protein group detection among the 8 demultiplexed isobaric labels. 3400 protein groups were detected in all samples and ~4000 protein groups identified in >=50% of the runs. **B)** Box plot highlighting the avered protein abundance of all proteins in each sample. The blue line indicates the average protein abundance of the least variable samples 113,116-119 and 121 **C)** PCA plot of all 6 samples with the highest average protein abundances. 113 is a clear outlier while 116, 117, and 118 group together. **D)** A heatmap of the top 100 most variably expressed proteins across the biological replicates, highlighting intra- and inter-condition variation. **E)** Volcanoplot shows roughly 800 DEPs that have a |log2FC| > 0.25 and FDR adjusted p-value < 0.05 highlighted in red. Green dots represent proteins that met the log2FC cutoff but not the pvalue cutoff. Gray dots represent the proteins that did not meet any cutoff. **F)** Venn Diagram of DEPs from rat mucosa proteome following 2wks of pBOO, DEPs of full thickness bladder following 2 weeks of pBOO and predicted early biomarkers of obstruction.

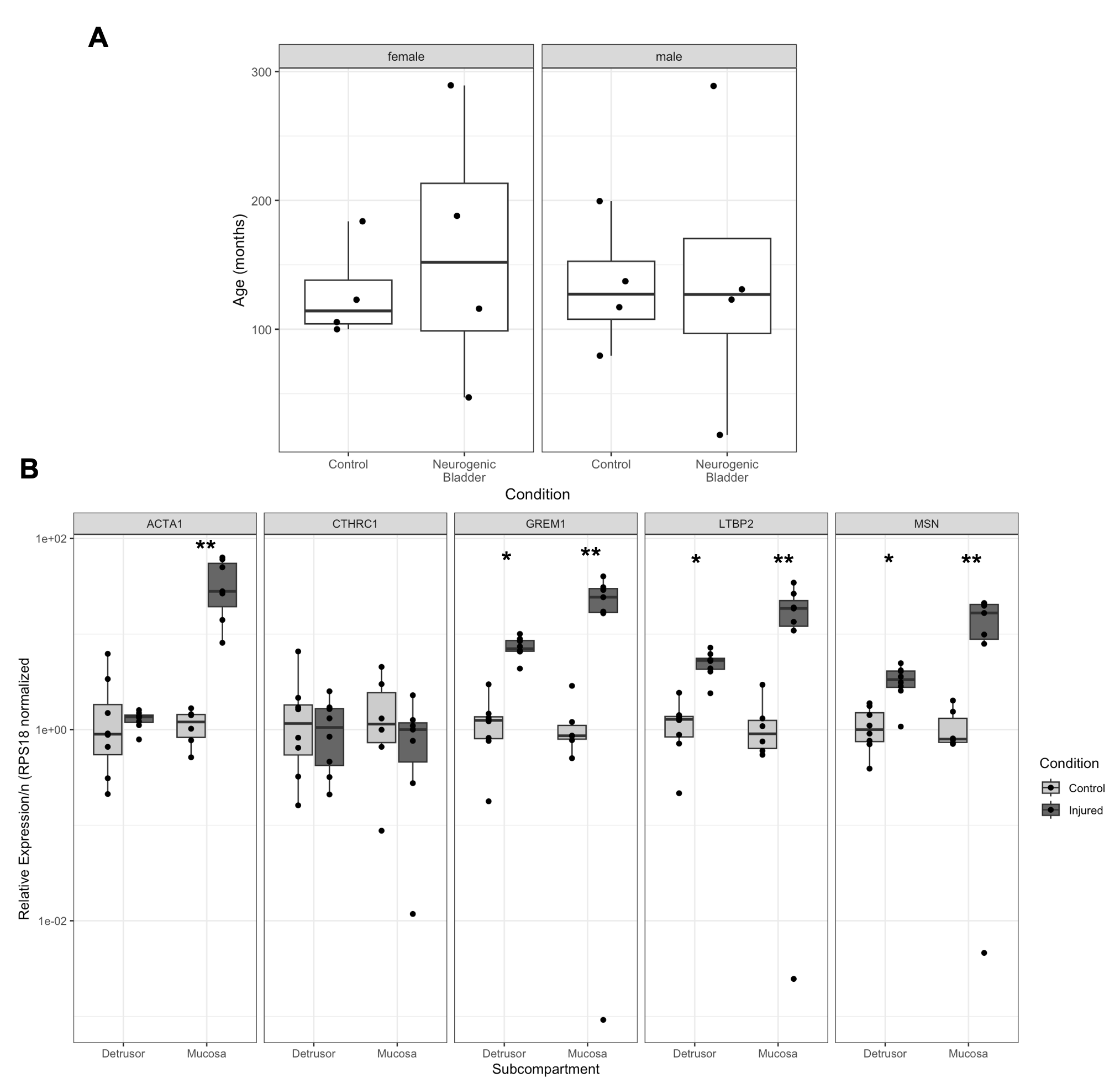

**Supplementary Figure 9. qPCR validation of predicted biomarker expression in human NGB**

**A)** A total of 8 male and female pediatric patients with Neurogenic bladder or control bladders were identified for **B)** qPCR analysis of expression of select biomarkers in separated Detrusor and Mucosa bladder compartments.

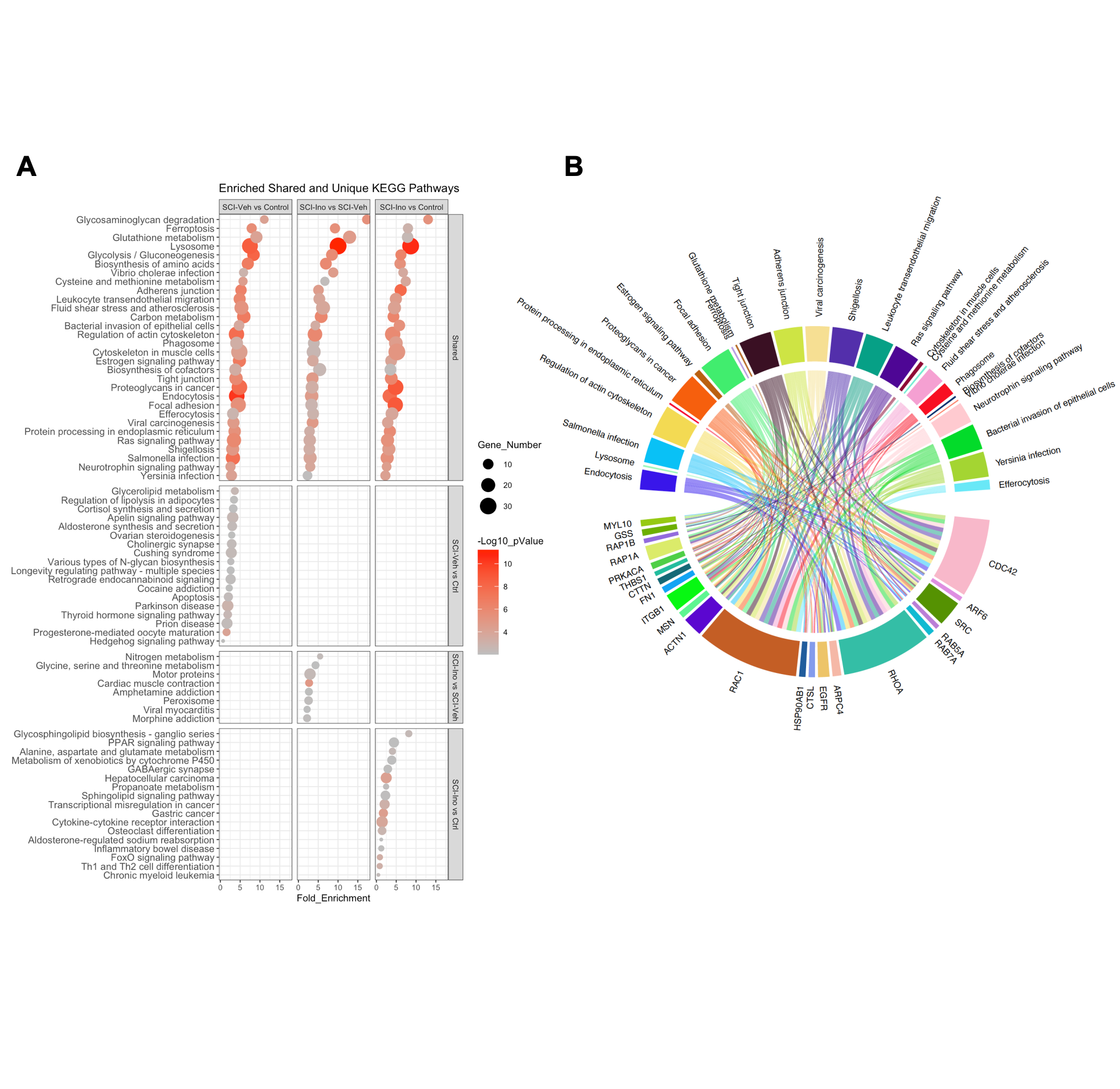

**Supplementary Figure 10. SCI and inosine-altered pathways and associated proteins analysis of urinary proteome**

Differentially expressed urinary proteins from three separate comparisons, i) spinal cord injury + vehicle vs. control (SCI-Veh vs Ctrl), ii) spinal cord injury + inosine vs. spinal cord injury + vehicle (SCI-Ino vs. SCI-Veh) and iii) SCI-Ino vs Ctrl were subjected to enrichment analysis of KEGG pathways. **A)** A dotplot of the enrichment results are shown in which pathways are depicted on the left-side y-axis, the extent of the pathway enrichment is on the x-axis, the right-side y-axis indicates whether a pathway is enriched in all comparisons (shared) or unique to one of the three comparisons. The size of the dots indicates the number of genes associated with that pathway. The color of the dots indicates the extent of statistical significance of the enrichment. **(B)** Chord diagram showing only the urinary DEPs associated with 4 or more of the shared enriched pathways.

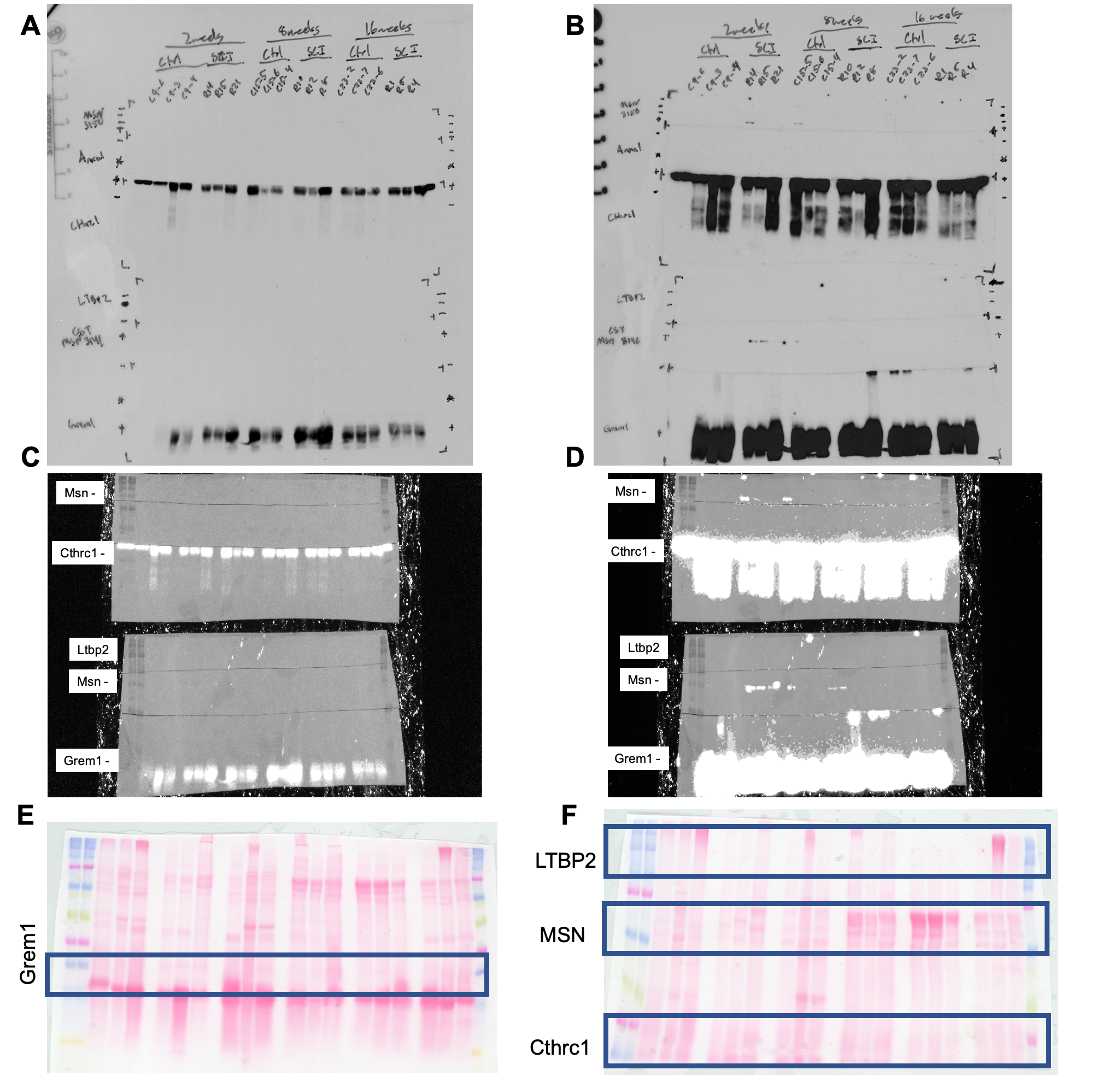

**Supplementary Figure 11. Uncropped immunoblots of rat urine samples**

Exposure of radiography film capturing signal from two membranes loaded with 50 µg of rat urine protein from control and spinal cord injured animals at 2, 8 and 16 weeks for 1 min **(A)** or 1 hr **(B)**. Digital signal capture from two membranes loaded with 50µg of rat urine protein from control and spinal cord injured animals at 2, 8 and 16 weeks for 1 min **(C)** or 1 hr **(D). (E & F)** Ponceau S signal from two membranes loaded with 50 µg of rat urine protein from control and spinal cord injured animals at 2, 8 and 16 weeks. Highlighted in blue boxes are the regions used to normalize the signal of each corresponding antibody

| **Target** | **Species** | **Forward Sequence** | **Reverse Sequence** |
| --- | --- | --- | --- |
| Cthrc1 | Rn | AGGGAAGTGGTAGACCTGTATAAT | TTGGGGTCCAGGATTCCTCA |
| Ltbp2 | Rn | GAAGGTACAAATGCAGCACCC | GGGGTTAGCACACTGTCCAT |
| Msn | Rn | CCGCAGTGAGGAAGAACGTA | GCTGTTTGGTATTGCCCTGC |
| Grem1 | Rn | AGTAGCGGCACTTTCCTTCG | TGCCGTGCGATTCATTCTGT |
| Gapdh | Rn | TCTCTGCTCCTCCCTGTTCT | CCGATACGGCCAAATCCGTT |
| Rps18 | Rn | ACAACGTGACTCCAGACATGA | GCCTTCTTTTGAGCAACCCG |
| Sdha | Rn | AGTCTATCGCTGAGGCTTGC | CGCGTCTGAAACTTTAGCGG |
| Bact | Rn | ACAACCTTCTTGCAGCTCCTC | CCTTCTGACCCATACCCACC |
| Acta1 | Rn | ACACCATGTGTGACGAGGAC | AGGGTCAGGATACCTCGCTT |
| CTHRC1 | Hs | CAGGACCTCTTCCCATTGAAGC | GCAACATCCACTAATCCAGCACC |
| LTBP2 | Hs | CTGCACAGATGACAACGAGTGTC | AGAGTGTAGCCAGGGTAGCAGA |
| MSN | Hs | CTGATGGAGAGGCTGAAGCAGA | ACGCTTCCGTTCCTGCTCAAGT |
| GREM1 | Hs | TCATCAACCGCTTCTGTTACGGC | CAGAAGGAGCAGGACTGAAAGG |
| GAPDH | Hs | GTCTCCTCTGACTTCAACAGCG | ACCACCCTGTTGCTGTAGCCAA |
| RPS18 | Hs | GCAGAATCCACGCCAGTACAAG | GCTTGTTGTCCAGACCATTGGC |
| SDHA | Hs | GAGATGTGGTGTCTCGGTCCAT | GCTGTCTCTGAAATGCCAGGCA |
| BACT | Hs | CACCATTGGCAATGAGCGGTTC | AGGTCTTTGCGGATGTCCACGT |
| Acta1 | Hs | AGGTCATCACCATCGGCAACGA | GCTGTTGTAGGTGGTCTCGTGA |

**Supplementary Tables**

**Supplementary Table 1.** qPCR primer sequences

**Supplementary Table 2.** Antibody information

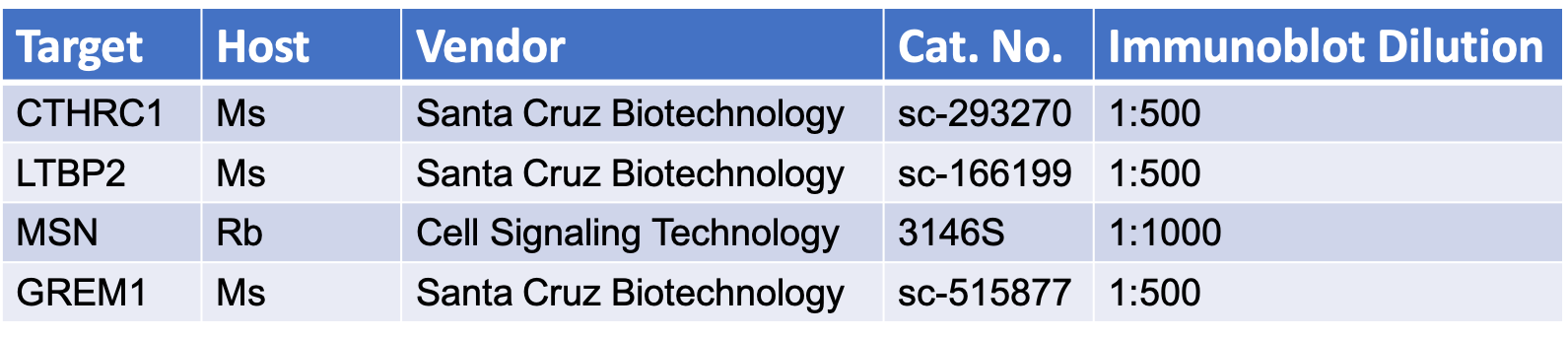

**Supplementary Table 3. 6 of 11 secreted biomarkers have no previous publications demonstrating urine detection in human samples.**

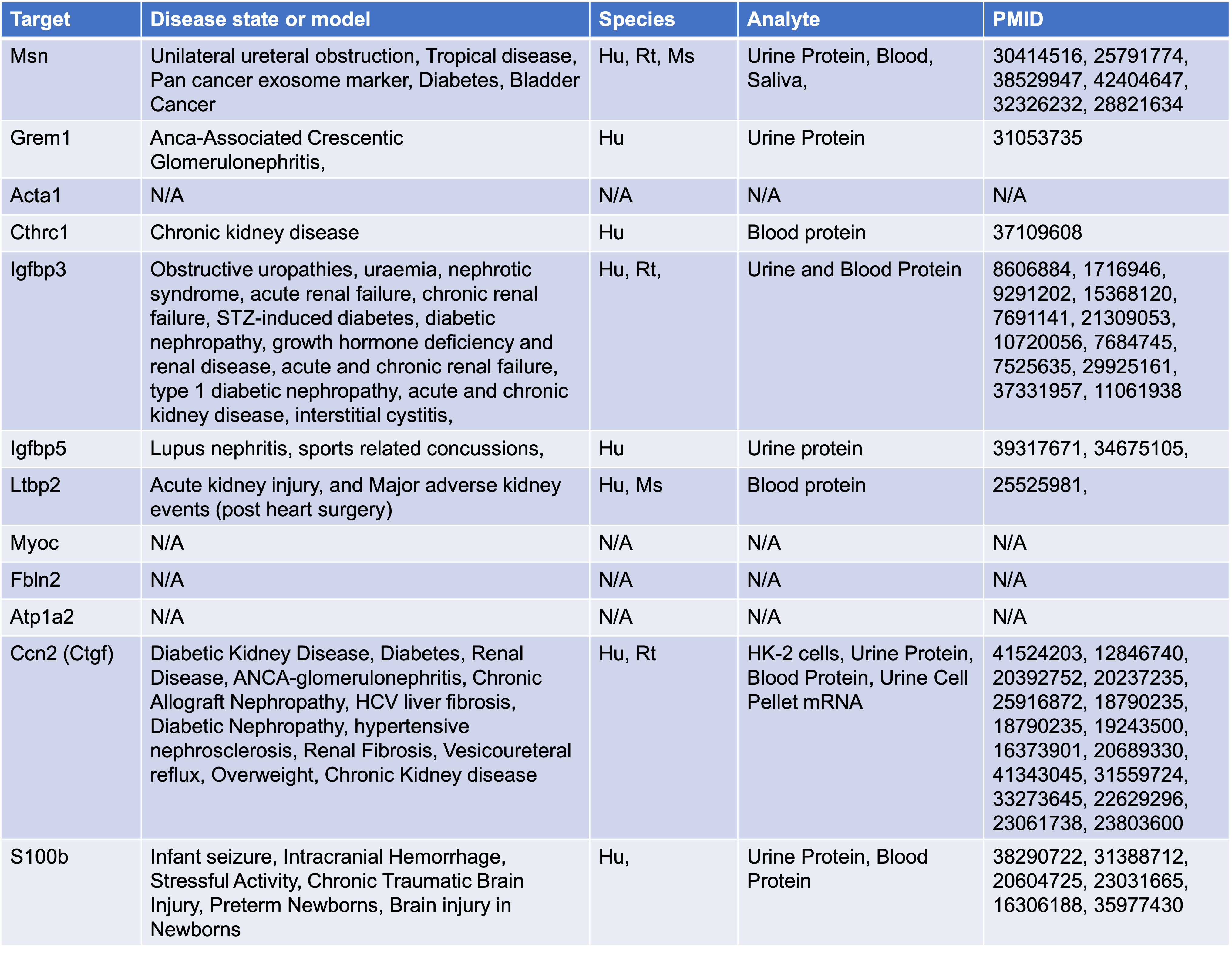
